# Germline maintenance through the multifaceted activities of GLH/Vasa in *Caenorhabditis elegans* P granules

**DOI:** 10.1101/663641

**Authors:** Elisabeth A. Marnik, J. Heath Fuqua, Catherine S. Sharp, Jesse D. Rochester, Emily L. Xu, Sarah E. Holbrook, Dustin L. Updike

## Abstract

Vasa is a highly conserved member of the ATP-dependent DEAD box helicase family, a multipotency factor, and a critical component for the specification and maintenance of the germline. Its homologs have been shown to regulate translation, small RNA amplification, and serve as a molecular solvent for single-stranded RNA; however, the function of Vasa’s defining domains and what they interact with are unclear. To address this, 28 mutant alleles of the *C. elegans* Vasa homolog GLH-1 were generated in conserved motifs. Mutations in the flanking and helicase domains show that GLH-1 retains its association with P granules through its helicase activity and not through static interactions with other P-granule proteins. Changes outside of these domains retain GLH-1 in P granules but still compromise fertility, and removal of glycine-rich repeats progressively diminish P-granule wetting-like interactions at the nuclear periphery. A mutation that facilitates Vasa aggregation was previously leveraged in insects and mammals to identify the transient association of Vasa with piRNA amplifying Argonautes. This same mutation in GLH-1 also stimulates aggregation and association with Argonautes, suggesting that the transient amplifying complex is evolutionarily conserved even though the method of piRNA amplification in *C. elegans* is not. Mass spectrometry analysis of proteins that co-immunoprecipitate with wild type and mutant GLH-1 reveal an affinity for all three PCI (26S <u>P</u>roteasome Lid, <u>C</u>OP9, eIF3) scaffolding complexes, which regulate protein turnover and translation, and a possible aversion for ribosomes and the 26S proteasome core. These results suggest that phase-separated P granules compartmentalize the cytoplasm to exclude large protein assemblies and emphasize the role of Vasa homologs in maintaining proteostasis.

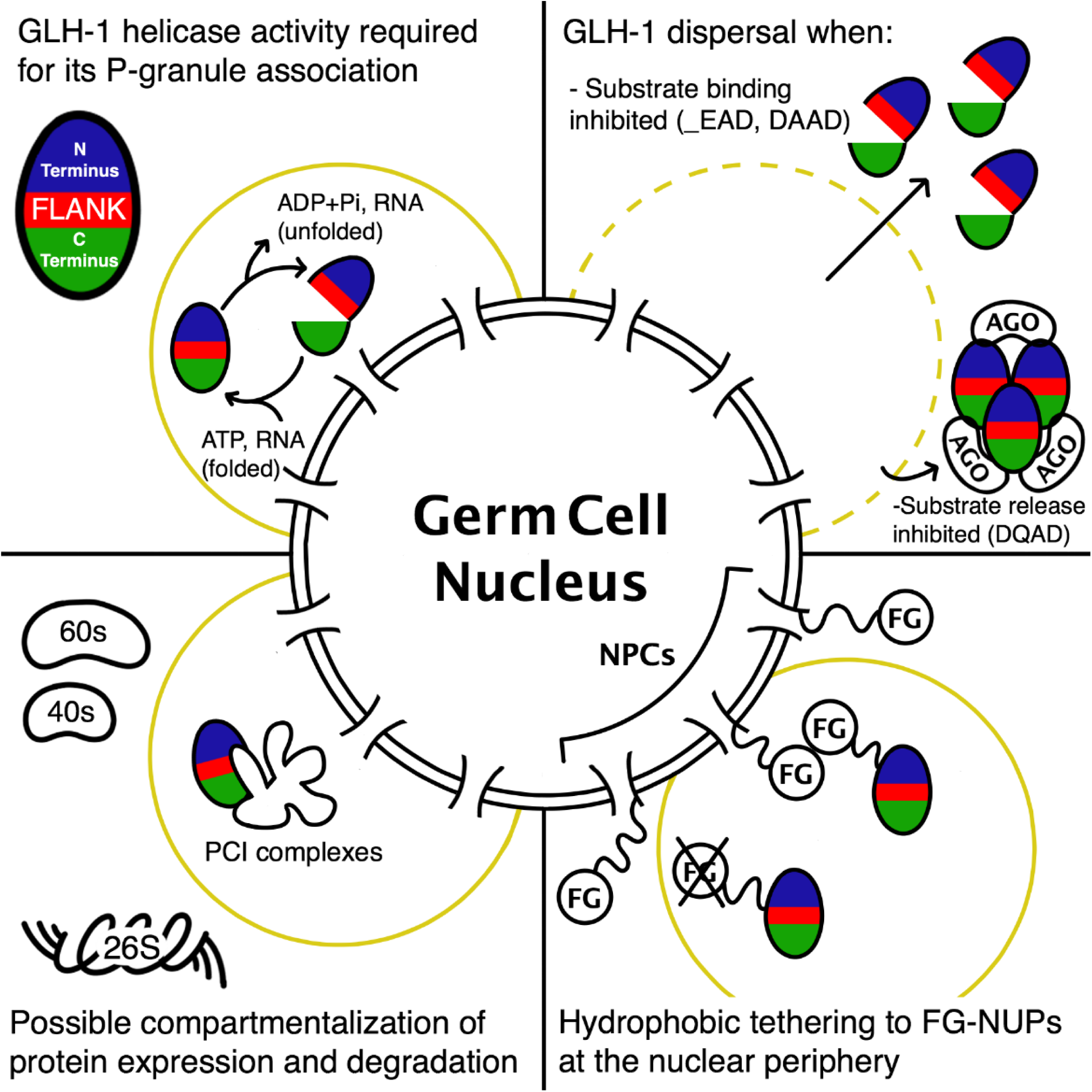
GRAPHICAL ABSTRACT.

**HIGHLIGHTS:**

- GLH-1/Vasa helicase activity is required for germ granule association and the flanking domain is critical component of this helicase activity.
- GLH-1 and GLH-2 glycine-rich FG-repeats increase the coverage or wetting-like properties of germ granules at the nuclear periphery.
- Locked GLH-1 helicase domains increase association with Argonaute proteins, resembling small RNA transient amplifying complexes observed in insects and mammals.
- GLH-1 has an affinity for all three PCI (26S Proteasome Lid, COP9, eIF3) scaffolding complexes, emphasizing a role in protein translation and turnover.

## INTRODUCTION

Germ cells and somatic cells from an individual carry identical copies of DNA; yet, only germ cells have the potential to give rise to all the cell types of each subsequent generation. This suggests that epigenetic factors confer a germ cell’s totipotent and immortal potential. These epigenetic factors are not limited to chromatin modifications, but also reside in the germ cell cytoplasm, or germ plasm. In some cases the presence of germ plasm alone has been sufficient to reprogram somatic nuclei to restore cellular potency and immortal potential (reviewed in Strome and Updike, 2015). Germ plasm contains a heterogeneous mix of RNA and protein that is not expressed in differentiating somatic tissue. In some animals these germ-cell specific ribonucleoproteins (RNPs) phase separate in the cytoplasm to form what are called germ granules (reviewed in Marnik and Updike, 2019). Depletion of germ granules in *C. elegans* causes sterility and germ to soma transformation, suggesting they contain the cytoplasmic components that preserve germ-cell totipotency (Knutson et al., 2017; Updike et al., 2014). A conserved protein that is consistently observed in the germ plasm and germ granules across species is collectively known as Vasa. Vasa and its homologs are required for germline specification, and have been shown more recently to influence somatic multipotency during development, regeneration, and tumorigenesis (reviewed in Poon et al., 2016). Therefore, understanding Vasa’s molecular function and its complex role as a multipotency factor is critical.

Vasa was cloned in Drosophila just over 30 years ago as a DEAD-box helicase with homology to the eukaryotic initiation factor-4A (eIF4A) (Hay et al., 1988; Lasko and Ashburner, 1988), and a binding partner to translation initiation factor (eIF5B) (Carrera et al., 2000). These findings strongly suggested that Vasa and its homologs function to initiate and/or regulate translation in the germline, which was subsequently demonstrated by the eIF5B-dependent accumulation of *Gurken* and *mei-P26* in Drosophila (Johnstone and Lasko, 2004; Liu et al., 2009). A more recent focus has been on Vasa’s RNA-independent interaction with Argonaute proteins through a transient amplifying complex that impacts ping-pong-mediated piRNA amplification (Dehghani and Lasko, 2016; Kuramochi-Miyagawa et al., 2010; Malone et al., 2009; Megosh et al., 2006; Wenda et al., 2017; Xiol et al., 2014). Other studies have demonstrated the ability of Vasa homologs (i.e. DDX4) to form phase separated organelles in cell culture that melt nucleic acid duplexes and act as a solvent for single-stranded RNA (Nott et al., 2015, 2016). The role of Vasa homologs in translational regulation, piRNA amplification, and as an mRNA solvent demonstrates the protein’s diversity of functions within the germline (reviewed in Lasko, 2013).

Phenotypes of various mutant Vasa alleles in Drosophila reflect this diversity of function. Strong alleles exhibit recessive female-sterility in homozygotes due to defective oocyte development. Moderate Vasa alleles produce oocytes but after fertilization the resulting embryos arrest with posterior patterning defects. Mutants rescued with Vasa transgene carrying weak alleles permit some embryos to hatch and develop into adults that exhibit a range of fertility defects (Dehghani and Lasko, 2015). Vasa phenotypes are also diverse across organisms. For example, Vasa mutations in Drosophila cause female-specific sterility whereas mutations in the Vasa homolog MVH cause male-specific sterility in mice (Wenda et al., 2017). Furthermore, while Vasa is conserved across metazoans, some animals such as *C. elegans* amplify piRNA silencing through RNA-dependent RNA polymerases (RdRPs) instead of the ping-pong method used by insects and mammals that lack RdRPs. Because of these differences, a comparative analysis of Vasa in different organisms is needed to determine conserved and divergent functions of germline maintenance and specification.

The comparison of Vasa function in model organisms has traditionally been limited by available mutants, making it difficult or impossible to gain insight on structural motifs from available alleles exhibiting a wide range of phenotypes. This was especially true in *C. elegans* where the function of one of its Vasa homologs, GLH-1, could only be inferred from a small handful of alleles that still made truncated proteins (Spike et al., 2008). However, with the advent of CRISPR, modifications to endogenous genes can be made to replicate informative alleles in conserved residues. Using this approach, over two dozen site-directed mutant alleles of *glh-1* were created in a strain where the endogenous gene carried a C-terminal GFP∷3xFLAG fusion. Each modification was then examined to determine its influence on fertility and embryonic viability in the context of its effect on GLH-1 expression and distribution in the embryonic and adult germline. These results emphasize the role of GLH-1’s helicase activity to maintain P-granule association and provide insight into the functional domains that distinguish Vasa proteins from the dozens of other DEAD-box helicases encoded in the *C. elegans* genome.

Vasa protein interactions may be very transient, making them difficult to detect. Previous DEAD-box helicase studies have utilized mutations within the DEAD motif that are thought to either inhibit substrate binding or lock in bound substrates, with the idea of capturing different interaction partners at distinctive, and often transient, enzymatic steps (Cruciat et al., 2013; Pause and Sonenberg, 1992; Xiol et al., 2014; Yang et al., 2014). In this report, IP LC-MS/MS was used to identify proteins with increased GLH-1 association and examine what happens to those associations in the substrate inhibited or locked states. These results suggest that GLH-1 associates with evolutionarily conserved PCI (26S **P**roteasome Lid, **C**OP9 signalosome, e**I**F3) scaffolding complexes or “zomes” to regulate protein translation and degradation. In the locked state, GLH-1 shows increased affinity for a handful of Argonaute proteins, suggesting that a form of the transient amplifying complex is conserved, but that GLH-1 is not limited to just piRNA amplification. This comparative approach represents a significant advance toward understanding how GLH-1/Vasa functions as a multipotency factor within and outside of the germline.

## RESULTS

GLH-1 is just one of several dozen proteins enriched in germ granules, also known as P granules in *C. elegans*, but like Vasa, it is part of a germ-granule protein core that is conserved across multicellular animals. GLH-1 is a constitutive P-granule protein, meaning that it associates with P granules at all stages of adult and embryonic development (Figure 1A). Other core germ-granule proteins include those carrying Tudor domains, and Argonaute proteins that bind small RNAs (Figure 1B). The function of this protein core in germ granules is both intriguing and elusive, but studies across systems have demonstrated a role for this core in regulating protein expression and small RNA biogenesis in a way that ensures germ cell integrity. GLH-1 has three close paralogs (GLH-2, GLH-3, and GLH-4), but only GLH-1 and GLH-2 contain all the domains that define Vasa proteins. Vasa-defining domains include a glycine-rich repeat domain, a flanking domain that wraps in between N- and C-terminal DEAD-box helicase domains, and a negatively charged domain that precedes a terminal tryptophan (Figure 1C, Figure S1A & B). A conserved zinc-knuckle domain can be found in most Vasa homologs, but has been lost several times throughout evolution (Gustafson and Wessel, 2010)(Figure S1C). GLH-1 mutations, including nulls, are fertile at the permissive temperature of 20°C, but are sterile at 26°C (Kuznicki et al., 2000). This temperature sensitive (ts)-sterile phenotype stems from redundancy with other GLHs, as both *glh-2 glh-1* and *glh-1; glh-4* double mutants are sterile at permissive temperatures and exhibit a severe reduction or no germ cells and little to no sperm (Spike et al., 2008, and this study). The *C. elegans* genome encodes about 50 DEAD-box helicases. Of these, the GLHs, RDE-12, VBH-1, LAF-1, DDX-19, and DDX-17 have glycine-rich repeats and (with the exception of DDX-17) have previously been shown to associate with P granules (Gruidl et al., 1996; Hubert and Anderson, 2009; Sheth et al., 2010; Shirayama et al., 2014); however, outside GLH-1 and GLH-2, none contain a full repertoire of Vasa domains (Figure 1C).

**Figure 1.**
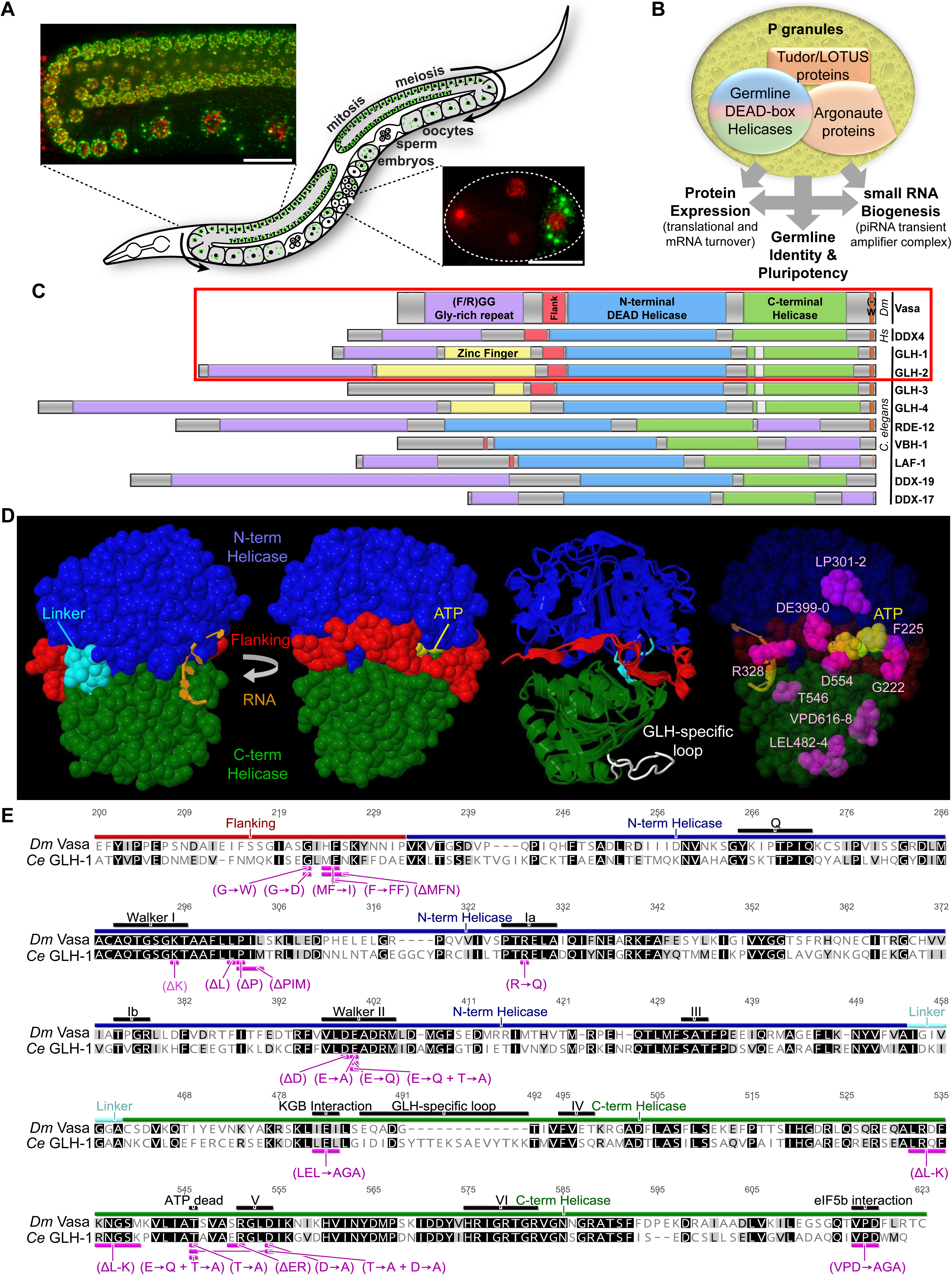
GLH Proteins in the *C. elegans* germline. A) GLH-1∷GFP∷3xFLAG in P granules and mCherry:His2B-marked chromatin. Inserts provide context for expression in the germline loop and 4-cell embryo. B) Schematic depicting the function of core proteins in P granules. C) Conservation of Vasa/DDX4-like DEAD box helicases in *C. elegans*. A red box surrounds proteins that contain all four Vasa-defining domains (glycine-rich in purple, flanking in red, N- and C-terminal helicase in blue and green, and negatively charged residues before a terminal tryptophan in orange). The GLH-specific loop is shown in white. D) Crystal structure of Vasa showing front and back views of the flanking and helicase domains in relation to ATP and RNA-binding pockets (as determined by Sengoku et al., 2006). Image 3 is an overlay of Vasa (ribbon) with an iTasser-predicted model of GLH-1 (backbone) that shows the location of the GLH-specific loop (white). Image 4 shows key amino acid residues targeted in this study and their location within the Vasa protein structure. E) Sequence alignment of the flanking and helicase domains in Drosophila Vasa with *C. elegans* GLH-1. Protein domains and mutations are indicated (purple). The K295A mutation was not obtained. The ΔER550-1 was not sustainable.

The structure of Drosophila’s Vasa flanking and helicase domains have been determined, showing that the N-(blue) and C-terminal (green) RecA-like DEAD-box domains interact upon RNA and ATP-binding (Figure 1D, Sengoku et al., 2006). ATP hydrolysis is coupled with RNA helicase activity, destabilizing RNA duplexes in a non-processive manner. The flanking domain (red) wraps around the side when the helicase domains are in the closed conformation. Because of the high conservation between Vasa in Drosophila and GLH-1 in *C. elegans* (Figure 1E), iTasser was used to model the structure of GLH-1 based on Vasa (Figure 1D, third image overlay). Except for a GLH-specific loop (white), the predicted structure is nearly identical. From this, several key residues and their relation to ATP and RNA-binding sites and helicase interphases can be identified (Figure 1D, fourth image).

New alleles of *glh-1* were initially obtained from an EMS mutagenesis screen for P-granule phenotypes. Unlike previous screens that utilized transgenes expressed from an array, this screen was performed in a strain where endogenous GLH-1 was tagged with GFP∷3xFLAG, allowing for the recovery of intragenic mutations. While most *glh-1* alleles from the screen attenuated GFP expression, one allele dispersed GLH-1∷GFP throughout the cytoplasm and resembled GLH-1 staining patterns previously associated with original *glh-1* alleles (Spike et al., 2008). Sequencing *glh-1* revealed a Gly to Asp (G→D) change (Vasa position 222) caused by a single base pair mutation within the flanking domain, suggesting that flanking domain function is required for GLH-1’s association with P granules (Figure 2A). Until now, the function of Vasa’s flanking domain was unknown because alleles within this domain did not previously exist. CRISPR was used to generate four additional alleles with mutations in the flanking domain, and all recapitulate the GLH-1∷GFP dispersal phenotype (Figure S2, Table S1). GLH-1-granule dispersal and expression intensity was quantified in ten gonad arms under fixed exposure conditions, as was the impact on embryonic lethality and fertility at permissive and restrictive temperatures (Figure S2, Figure 3). For comparison, a complete *glh-1* deletion (*Δglh-1*) that expresses only GFP∷3xFLAG, and a *glh-1* transcriptional reporter (GFP∷3xFLAG separated from *glh-1* with an intercistronic rSL2 spacer, *glh-1* Txn GFP, Figure 2A) were generated from the parental strain. Interestingly, GFP expression in the deletion is about three times as bright, suggesting that GLH-1 protein negatively autoregulates its own expression. Fertility defects at the restrictive temperature for flanking domain mutations is comparable to the *glh-1* deletion, showing that these flanking domain mutations reflect the phenotype of null alleles.

**Figure 2.**
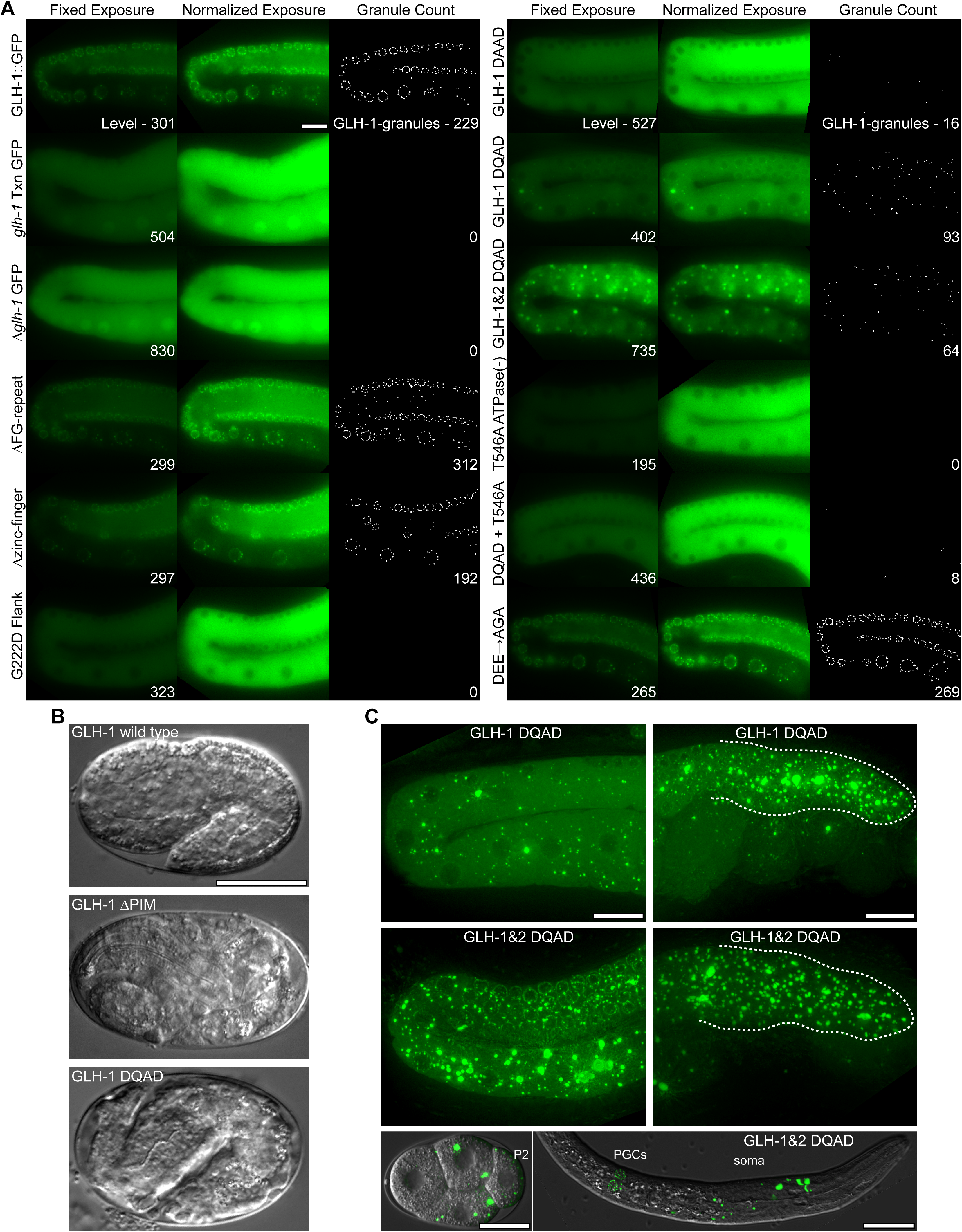
Level and distribution of mutant forms of GLH-1 in *C. elegans*. A) GLH-1∷GFP∷3xFLAG expression in the germline loop. Fixed exposures (left) were normalized (middle) to better view the distribution of fluorescence. GLH-1 granules were quantified using ImageJ (right). In ΔPIM and DQAD mutants, embryos arrest in the elongation phase. C) GLH-1(DQAD)∷GFP∷3xFLAG accumulation in proximal (left) and distal (right) germlines is enhanced in GLH-2(DQAD). In double mutants, GLH-1(DQAD) aggregates persist in somatic blastomeres and the soma of hatched worms (bottom).

To see if compromising GLH-1 helicase activity also caused its dispersal, 12 additional strains were created that either replicated canonical Drosophila alleles of Vasa in endogenous GLH-1∷GFP∷3xFLAG or introduced changes in key conserved residues (Figure 1E)(Dehghani and Lasko, 2015, 2016; Sengoku et al., 2006). Attempts to generate a K→A mutation in the Walker I motif (Vasa position 295) to knock out helicase activity were unsuccessful but yielded three mutations (ΔL, ΔP, ΔPIM) immediately to the right of the Walker I motif that all had the dispersed GLH-1∷GFP phenotype (Figure 3, Figure S2). Subsequently, a T→A mutation was generated just before motif V (Vasa position 546) that had previously been shown to abolish the ATPase activity of Vasa in vitro (Sengoku et al., 2006), and this allele disperses GLH-1 and causes fertility defects at the restrictive temperature (Figure 2A, Figure 3). The DEAD-box in motif II is also essential for ATP hydrolysis (Pause and Sonenberg, 1992), so a deletion of the aspartic acid (D) from the DEAD-box (**_**EAD) was generated (Vasa position 399) as was an E to A (D**A**AD) substitution (Vasa position 400), both of which disperse GLH-1 with ts-fertility defects (Figure 2A, Figure 3). When the **_**EAD deletion and the D**A**AD substitution are put into both *glh-1* and *glh-2* to create double mutants it results in fertility defects at both permissive and restrictive temperatures (Figure 3). When taken together these results suggest that 1) helicase activity is required for fertility, 2) GLH-1 associates with germ granules by virtue of its helicase activity and not through its structural motifs, that 3) the flanking domain is integral to the helicase activity, and that 4) helicase activity is not required for GLH-1 to negatively autoregulate its expression.

**Figure 3.**
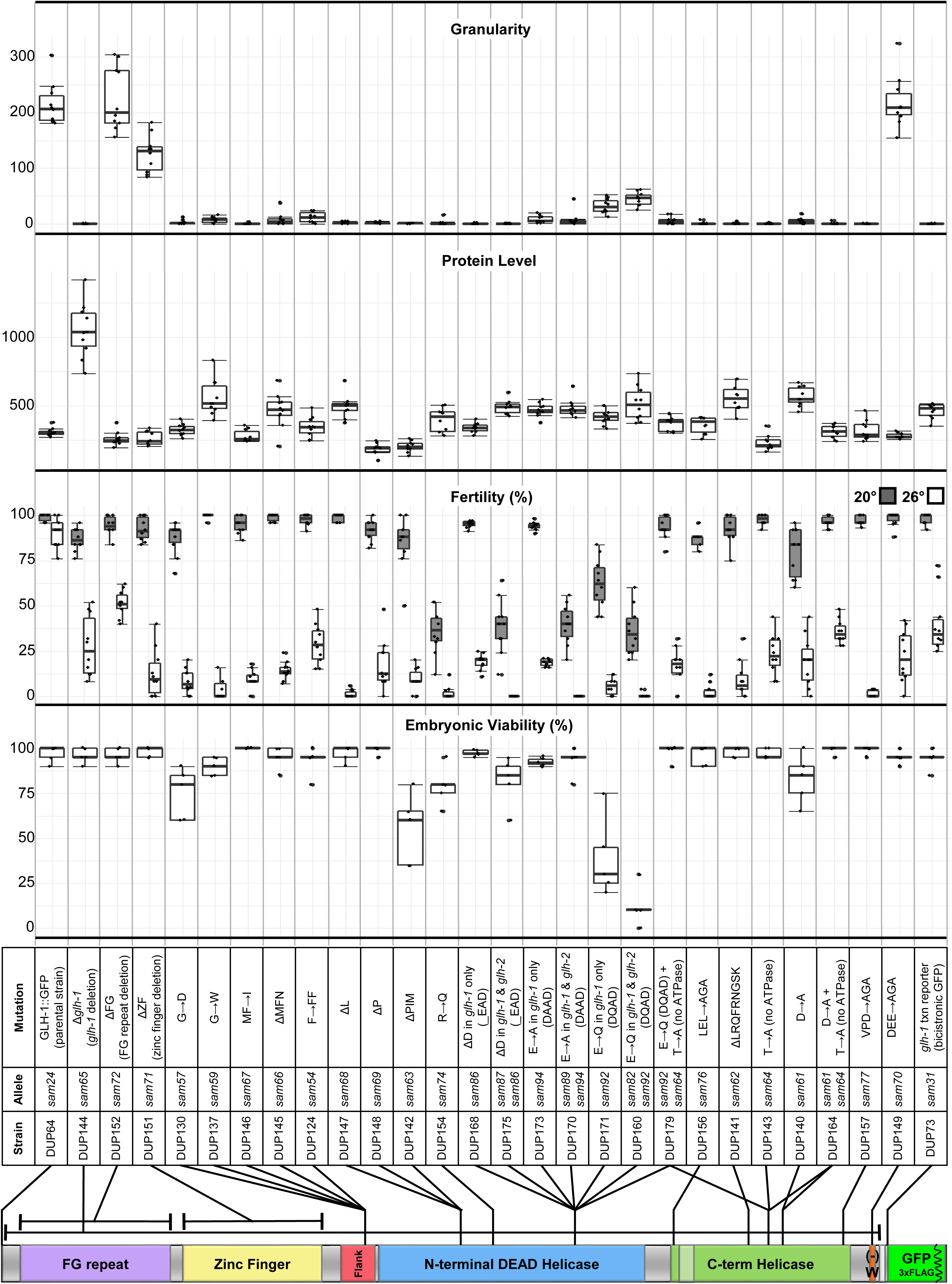
Consequences of GLH-1 mutant alleles. From top to bottom: Comparison of granularity (GLH-1 granules), GLH-1 protein level, fertility at permissive (20°C) and restrictive (26°C) temperatures, and embryonic viability in GLH-1 mutants. Mutation details, strain and allele names and their respective locations are indicated.

Vasa was identified as a component of a transient Amplifier complex that mediates piRNA amplification in what is called the ping-pong loop (Wenda et al., 2017; Xiol et al., 2014). This association was detected using a Vasa D**Q**AD mutation (position 400) that is thought to prevent the release of ATP hydrolysis products, facilitating the accumulation of larger Vasa-containing aggregates. While *C. elegans* use a ping-pong independent method to amplify piRNA mediated silencing, D**Q**AD substitutions were introduced into *glh-1* single and into *glh-1* and *glh-2* double mutants to see if similar large aggregates form (Figure 2A). Unlike **_**EAD and D**A**AD, the D**Q**AD substitution is more severe than the *glh-1* deletion, meaning less fertile at 20°C and most of its embryos arrest during elongation (Figure 2B, Figure 3). Also, instead of being completely dispersed in the cytoplasm like **_**EAD and D**A**AD, D**Q**AD causes some GLH-1 to accumulate in large cytoplasmic aggregates, primarily in the shared cytoplasm of the distal germline (Figure 2C). Embryonic lethality and large aggregate formation of GLH-1(D**Q**AD) becomes more pronounced when the D**Q**AD substitution is also introduced into GLH-2. In the double mutant, large GLH-1∷GFP(D**Q**AD) aggregates are no longer cleared from somatic blastomeres, and some persist in various somatic cells during larval development (Figure 2C). These double mutants can be passaged for only a couple generations and must be maintained over a balancer. It should be noted that while these large aggregates have been ascribed as a specific transient state, their observation in D**Q**AD but not in **_**EAD or D**A**AD introduces the possibility that D**Q**AD creates a neomorphic allele prone to unspecific germline aggregate formation. The impact of the D**Q**AD mutant has not been thoroughly characterized in vivo, and existing data from in vitro experiments are not enough to understand why **_**EAD and D**A**AD disperses GLH-1 or Vasa while D**Q**AD causes them to aggregate. A prevailing assumption is that mutations like **_**EAD and D**A**AD inhibit the binding of ATP and RNA substrates while in D**Q**AD these substrates are bound but not released, but further characterization of these mutations is needed, both in vitro and in vivo

GLH-1 has been positioned upstream of PGL proteins in the embryonic P-granule assembly pathway, but in adult germlines the association is more mutualistic (Hanazawa et al., 2011; Kawasaki et al., 2004; Kuznicki et al., 2000; Updike et al., 2011). Both PGL-1 and GLH-1 colocalize at all stages of development in wild-type animals, except for a brief resurgence of small somatic PGL-1 granules around the 30-50 cell stage of embryogenesis (Figure 4A). While PGL-1 is dispersed in zygotes of the GLH-1(G222D) mutant, it reassembles into P granules despite disperse GLH-1 beginning at the 4-cell stage and largely stays associated with P granules in the adult germline (Figure 4B). A similar pattern is observed with mCherry-tagged PRG-1, the PIWI Argonaute in *C. elegans*, which maintains its association with P granules in the adult germline in *Δglh-1* and GLH-1(**_**EAD) mutants (Figure 4D). In contrast, the large GLH-1 aggregates in D**Q**AD mutants contain both PGL-1 and PRG-1, suggesting that other P-granule components are locked in these large GLH-1(D**Q**AD) aggregates (Figure 4C,D).

**Figure 4.**
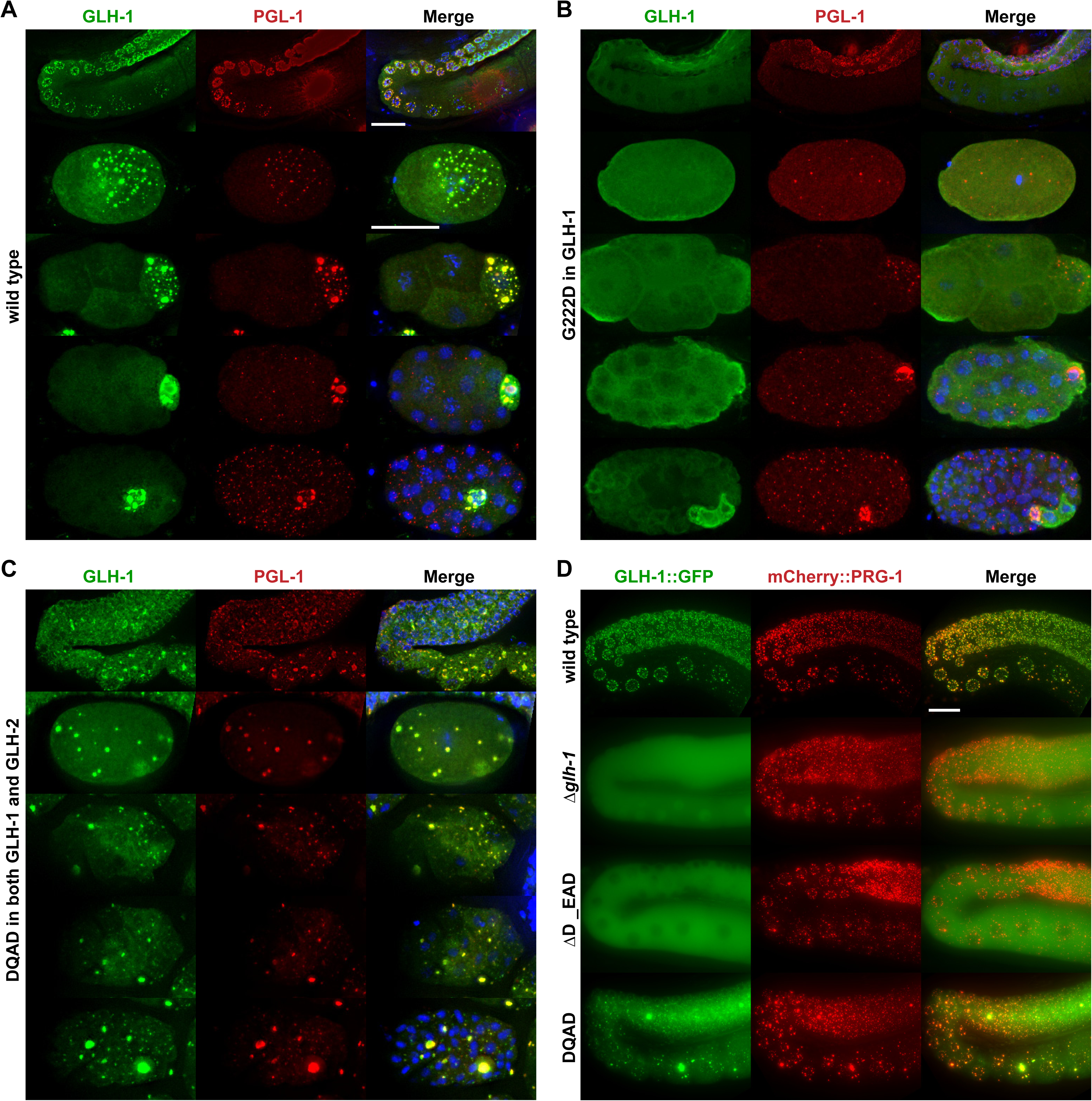
Colocalization of P-granule components in wild-type and GLH-1 mutants. A-C) Immunostaining of GLH-1 (green) and PGL-1 (red) in fixed germlines and 1, 4, 16, and 32-cell embryos. D) GLH-1∷GFP and mCherry∷PRG-1 in the germline of living worms.

It is possible that the GLH-1(D**A**AD) mutation inhibits ATP binding or hydrolysis, maintaining the N- and C-terminal domains in their open configuration, while the D**Q**AD mutation remains bound to hydrolyzed ATP with N- and C-terminal domains closed, as has been proposed for Vasa. In that case, an indirect assessment would be to inhibit ATP hydrolysis to see if it suppresses the embryonic lethality and fertility defects of D**Q**AD. To assess this possibility, CRISPR was used to generate the (D**Q**AD) + (T to A) (Vasa position 546) double mutant. This strain no longer exhibits large GLH-1 aggregates (Figure 2A), and the fertility defects and embryonic lethality are suppressed at the permissive temperature (Figure 3). These results support the idea that D**Q**AD aggregates are locked in a closed-conformation transient state. Since this transient state in other systems is associated with piRNA amplification (Dehghani and Lasko, 2016; Wenda et al., 2017; Xiol et al., 2014), TaqMan-based small-RNA expression assays were used to address whether GLH-1 impacts the accumulation of 21UR-1 abundance in *glh-1* mutants. 21UR-1 does not accumulate in a *prg-1* mutant when compared to wild-type, and the *glh-1* deletion and (**_**EAD) deletion show a small but insignificant reduction in 21UR-1 levels (Figure S3A). 21UR-1 levels in the D**Q**AD allele and the allele substituting the negative charge preceding the terminal tryptophan surprisingly show increases in 21UR-1 levels, but with large amounts of variability that render them insignificant. This is a departure from the effect of identical Vasa alleles in insects that reduce piRNA accumulation and may suggest that GLH-1 interacts with multiple Argonautes to affect a variety of small RNAs, including microRNAs used to normalize the TaqMan-based expression assay. Small RNA sequencing will be required to determine the effects of these *glh-1* alleles.

One Vasa mutation shown to uncouple ATP hydrolysis from its helicase activity in vitro is D to A (Vasa position 554), which lies at the interphase of Vasa’s N- and C-terminal helicase domains (Sengoku et al., 2006). This mutation also has a mild dominant negative phenotype in *C. elegans*, showing increased embryonic lethality and fertility defects (Figure 3). To determine if this could be caused by GLH-1 expending ATP but not coupling it with helicase activity, the analogous T546A was introduced to inhibit ATPase activity (Figure S2). This double mutant suppressed both the embryonic and fertility defects of the D554A mutant (Figure 3). To further test this idea, an R to Q mutation (Vasa position 328) was engineered to disrupt helicase activity in the RNA-binding pocket with minimal impact on helicase structure – potentially uncoupling helicase activity from ATP hydrolysis. Like D554A, R328Q alleles also enhanced embryonic lethality and fertility defects (Figure 3). These alleles may suggest that expenditure of ATP uncoupled from helicase activity drives dominant Vasa and GLH-1 phenotypes. Two additional C-terminal helicase alleles were created to disrupt previously reported binding sites for KGB-1 (LEL→AGA) and eIF5b (VPD→AGA), however both dispersed GLH-1 and look like other helicase mutations (Figure 3).

Outside of the flanking and helicase domains are three Vasa-specific motifs: a glycine-rich FG-repeat, a zinc-knuckle/finger, and a terminal tryptophan immediately preceded by three negatively charged amino acids. Unlike mutations in the flanking and helicase domains, deletions and substitutions in these motifs have no or very little effect on GLH-1’s association with P granules in the adult germline (Figure 2A); however, each show compromised fertility at the restrictive temperature (Figure 3). While this demonstrates that GLH-1 function can still be impaired despite showing proper P-granule localization, it came as a surprise for the ΔFG-repeat strain since it was previously shown to facilitate contact with the nuclear periphery when ectopically expressed (Updike et al., 2011). Vasa proteins contain these glycine-rich repeats that are interspersed with either arginines or phenylalanines (reviewed in Marnik and Updike, 2019). These are intrinsically disordered motifs, and in the case of the FG-repeats of GLH, the interspersed phenylalanines form hydrophobic tethers with FG-repeats in the nuclear pore complex (NPC) to maintain a wetting-like appearance on the nuclear periphery (Figure 5). Unlike the adult germline, deletion of the FG-repeat in embryos causes larger GLH-1∷GFP granules in primordial germ cells and their precursors. Additionally, deleting the FG-repeat of GLH-2 in this background further increases the size of these granules (Figure 5A & B). Moreover, GLH-1 granules in these double mutants appear more spherical, suggesting that they are losing contacts that adhere them to the nuclear periphery. One potential role for FG-repeat tethering is to maximize coverage of NPCs to capture nascent transcripts exiting the nucleus. Another is to ensure the symmetric distribution of P granules as the P4 precursor divides into the two primordial germ cells, but no evidence supporting this has yet been observed in the double mutant. It’s possible the FG-repeats found in GLH-4, RDE-12, and DDX-19 could be functioning redundantly and masking an asymmetric distribution phenotype when the FG-repeat is deleted in GLH-1 and GLH-2.

**Figure 5.**
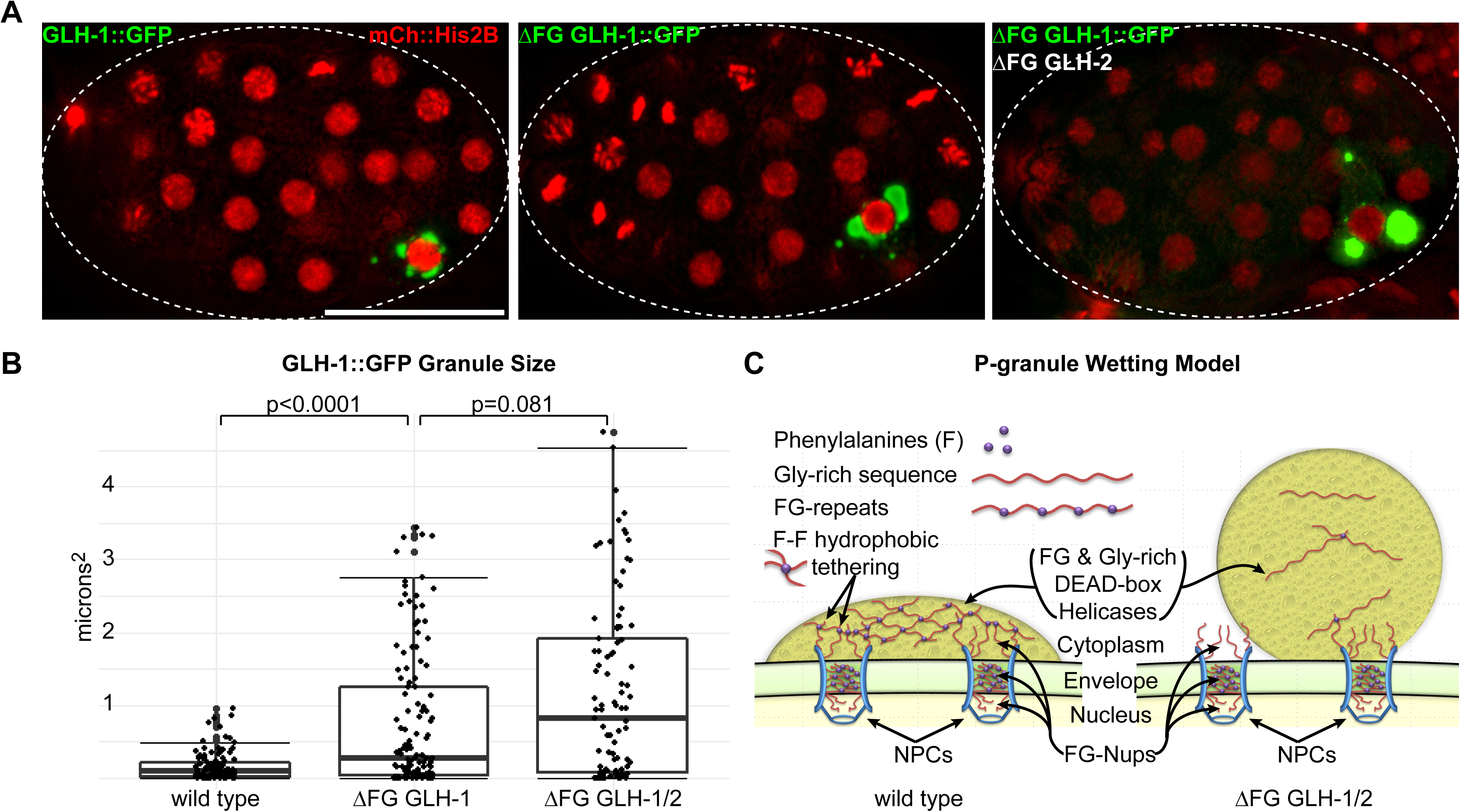
Glycine-rich FG repeats tether GLH-1 to the nuclear periphery. A) Distribution of GLH-1∷GFP (green) in the germline blastomere P4 of a wild-type embryo (left), embryo lacking FG repeats in GLH-1 (middle), and embryo lacking FG repeats in both GLH-1 and GLH-2 (right). mCherry∷H2B (red) marks chromatin. B) GLH-1 granule sizes in the germline blastomere P4 quantified using ImageJ. C) Model depicting the hydrophobic tethering of P-granule FG-repeat domains to FG-Nups at the nuclear periphery.

To get an idea of which proteins are loosely associated with GLH-1, and how these associations change when enzymatic activity is compromised in D**Q**AD and D**A**AD mutants, GLH-1∷GFP∷3xFLAG was immunoprecipitated with anti-FLAG agarose beads and replicates were submitted for LC-MS/MS analysis (Figure 6, Figure S3B). Pairwise comparisons between the *glh-1* transcriptional reporter driving GFP∷3xFLAG alone identified GLH-1 enriched proteins (Figure S2). As a proof of principle, nuclear pore complex (NPC) proteins and transport factors were identified among the 2505 proteins from the LC-MS/MS analysis (Figure 6, left column, blue, Figure S3). NPCs facilitate the interaction of P granules at the nuclear periphery, and several of them when targeted by RNAi cause P granules to detach and distribute in the cytoplasm (Updike and Strome, 2009; Voronina and Seydoux, 2010). On average, NPCs are enriched in the GLH-1 IP, and this enrichment shows a significant decrease (left shift) in both the D**A**AD and D**Q**AD mutants as would be expected with the dispersal of GLH-1 from the nuclear periphery in the mutants, confirming the robustness of the GLH-1 IP (Figure 6).

**Figure 6.**
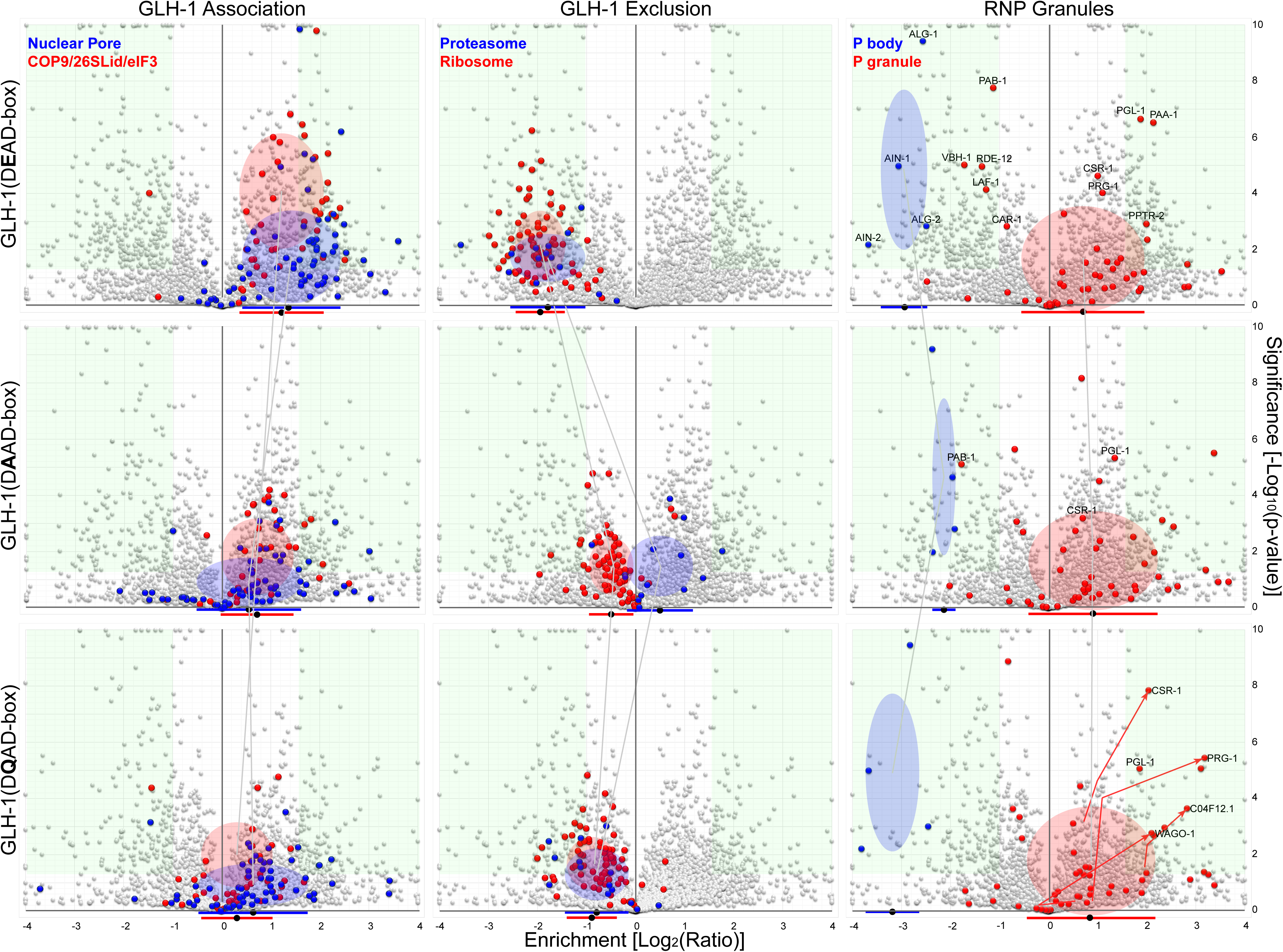
GLH-1 protein associations. Volcano plots show the significance and enrichment of proteins that immunoprecipicate with GLH-1∷GFP∷3xFLAG over the *glh-1* transcriptional reporter expressing GFP∷3xFLAG alone, as identified by LC/MS/MS. Left column: protein families showing an enriched GLH-1 association include nuclear pore proteins (blue) and subunits of PCI scaffolding complexes (26S Proteasome Lid, COP9 Signalosome, and eIF3). These associations decrease in DAAD (middle row) and DQAD (bottom row) mutants. Red and blue bars under the x-axis indicate the median and one standard deviation, and colored ovals do the same but also indicate the distribution of significance. Green boxes show normalized enrichment and exclusion greater than 2.5-fold and a p value < 0.05. Nuclear pore genes used in this analysis are indicated in Table S3. Middle column: protein families showing stronger enrichment for GFP∷3xFLAG alone include 20S core subunits of the 26S proteasome (blue) and subunits of the ribosome (red). Right column: proteins associated with RNP granules. P-body proteins (blue) and P-granule proteins (red).

Gene ontology was examined in the enriched subsets (>2.5-fold normalized increase, p value<0.05, Table S2), which identified most subunits of three evolutionarily conserved, multi-lobed scaffolding complexes collectively known as PCI complexes or “zomes” (Li et al., 2017). These include the COP9 signalosome, the regulatory Lid complex of the 26S proteasome, and the eIF3 translational initiation complex (Figure 6, left column, red; Table S3). One subunit of the COP9 signalosome called CSN-5 was previously identified through a yeast two-hybrid screen with GLH-1 as bait, and the interaction was confirmed through pull downs (Smith et al., 2002). Vasa-GST pull downs later confirmed the direct interaction with CSN5 is protective and evolutionarily conserved (Orsborn et al., 2007). This LC-MS/MS analysis suggests the structural conservation of all three “zomes” can facilitate increased association with GLH-1, and that this interaction is partially compromised in D**A**AD and D**Q**AD mutants.

Both COP9 and lid complexes modulate protein degradation by 26S proteasome through de-neddylation and deubiquitination, respectively (reviewed in Meister et al., 2016). Interestingly, subunits of the 20S core of the 26S proteasome are depleted in the GLH-1 IP, and this depletion is dampened (right shift) as GLH-1 becomes dispersed in the cytoplasm of D**A**AD and D**Q**AD mutants (Figure 6, middle column, blue). Whether GLH-1 is 1) sequestering these regulatory PCI complexes in P granules and away from the 20S proteasome core to antagonize protein degradation, 2) associating with COP9 and Lid complexes prior to degradation in somatic blastomeres, or 3) facilitating the cycling of cullin-RING E3 ubiquitin ligase (CRL) activity still needs to be determined. Degradation of P granules in somatic blastomeres is mediated by CRL activity with the CCCH-finger-binding protein ZIF-1 acting as a receptor (DeRenzo et al., 2003; Oldenbroek et al., 2012). RNAi depletion of transcripts encoding multiple 20S core proteasome subunits, regulatory lid subunits, and ubiquitins cause P-granule accumulation throughout the soma of arrested embryos (Updike and Strome, 2009). Interestingly, Drosophila Vasa is also regulated through CRL activity by two CRL-specificity receptors (Gus and Fsn) that compete for a single binding site on Vasa; the Gus receptor acts to stabilize Vasa and protect it from Fsn-mediated destabilization (Kugler et al., 2010). Gus and Fsn homologs were not enriched in our GLH-1 IP LC/MS/MS analysis, and the Gus binding sites of Vasa do not appear conserved in GLHs; instead, in its place are ancestral CCHC zinc-knuckle motifs that have been independently lost in insects, tardigrades, vertebrates, and some sponges and flatworms (Figure S1C). While little is known about this motif in GLH-1, some evidence suggests that zinc-knuckles may facilitate an interaction with an F-box containing P-granule protein called PAN-1 (Gao et al., 2012). An intriguing possibility is that insects developed a convergent method using Gus and Fsn to protect Vasa from proteasome degradation, and that COP9 and Lid regulatory subunit sequestration by GLH-1 has a similar protective effect.

P granules may also act to exclude 40s and 60s ribosomes, whose proteins, like those of the 20S proteasome core, are depleted in the GLH-1 IP (Figure 6, middle column, red). Again, this depletion is dampened (right shift) as GLH-1 becomes dispersed in the cytoplasm of D**A**AD and D**Q**AD mutants. This could suggest that 1) GLH-1 facilitates mRNA loading into eIF3 within P granules, priming them for ribosome assembly and translational expression as they exit P granules, or that 2) GLH-1 sequesters the eIF3 translation initiation complex from ribosomes as a general inhibitor of translation, or that 3) GLH-1 specifically inhibits the translation of P-granule enriched transcripts by associating them with eIF3 but protecting them from ribosome assembly. The increased association of GLH-1 with eIF3 refines proposed models of P-granule function in the germline and lays the ground work for follow up studies to determine whether the association between GLH-1 and the eIF3 complex mediates a positive or negative effect on translation.

Finally, several established P-body (blue) and P-granule (red) components were examined in the context of GLH-1 association or exclusion (Figure 6, right column, Table S3). Generally, the average dispersal of P-granule components changes very little in the D**A**AD and D**Q**AD mutants. The D**Q**AD mutant has been utilized in other systems to capture factors that associate transiently in the piRNA amplifier complex (Wenda et al., 2017; Xiol et al., 2014). Proteomics data from the D**Q**AD mutant were examined to see if any proteins increase in significance and association. Only four proteins showed this up-and-to-the-right shift from wild-type (DEAD) (red arrows), and they include the Argonaute proteins CSR-1, PRG-1, C04F12.1, and WAGO-1. This increased association of GLH-1 with Argonautes likely supports a complex similar to the piRNA transient amplifying complex described in insects and vertebrates, but one that is more broadly implicated in small RNA biogenesis and amplification and not specifically limited to piRNAs.

## DISCUSSION

The role of germ granules in inducing or maintaining germ-cell potency may come down to the molecular function of its individual components. In this study, a comparative structure-function analysis was performed on the *C. elegans* Vasa homologs. These results show that GLH-1’s helicase activity is necessary to maintain its tight association with P granules. Every edit of a conserved residue within the helicase domains caused GLH-1 to detach from P granules and disperse in the cytoplasm. Flanking domain mutations phenocopy this dispersal, suggesting that the flanking domain facilitates this helicase activity as it wraps between the N- and C-terminal RecA domains. Whether this means that GLH-1 localization is mediated through continuous unwinding of RNA substrates or continuous cycling of other protein interactions is unclear. However, GLH-1’s P-granule association is not mediated by its glycine-rich IDR, zinc-knuckle, negatively charged C-terminus or any inherent structural features on their own. Interestingly, while mutations in these domains do not disperse GLH-1 protein, they still exhibit close to the same degree of fertility defects at the restrictive temperature as the *glh-1* deletion, demonstrating one reason why these Vasa-defining domains have been conserved throughout evolution. The specific contribution of each of these domains will need to be ascertained through complimentary approaches that include 1) generating similar edits and deletions in paralogs to observe additive effects, as was done with the FG-repeat deletion in GLH-1 and GLH-2, and 2) by GLH-1 IP and LC/MS/MS in these mutant strains to determine which protein enrichments are lost and to find candidates to test for direct GLH-1 domain interactions.

One outstanding question is whether GLH-1 and P granules demonstrate an affinity to specific germline-expressed transcripts. Multiple attempts to immunoprecipitate and sequence RNA substrates of GLH-1 and PGL-1 under varying conditions have been performed by our group; however, follow-up single-molecule FISH studies have not demonstrated consistent P-granule enrichment of these identified substrates. These negative results likely reflect the non-sequence specific and transient manner in which core P-granule components like PGL-1 and GLH-1 interact with RNA, and they add weight to the idea that GLH-1 and Vasa proteins simply function as mRNA solvents in phase-separated P granules (Nott et al., 2016). This may eventually be resolved as RIP- and CLIP-seq technologies improve or the right P-granule target protein is found, but the idea that P-granules contain solvents to keep transcripts unfolded and accessible for sequence scanning by small RNA-bound Argonautes or other RNA-binding proteins is highly likely.

Dominant phenotypes were observed in R328Q and D554A mutations thought to uncouple ATP hydrolysis from RNA unwinding, which are possibly caused by increased energy expenditure that isn’t translated into enzymatic helicase activity. The D**Q**AD mutation is also dominant, but the D**A**AD mutation is not. This E to Q change may induce sterility because it accumulates large aggregates that sequester components from their normal function within P granules, or because they fail to dissolve in the soma. Since these aggregates persist in somatic blastomeres, the extent to which they resemble P granules or retain normal P-granule function is unclear. Therefore, caution should be maintained when interpreting whether D**Q**AD aggregates are capturing a transient amplifying complex or a novel aggregate altogether. Given that these dominant alleles are suppressed with an intragenic T546A mutation, they are likely anti- or neomorphic alleles. Another unreported deletion of ER residues in Motif V (Vasa position 550-551) caused a stronger dominant phenotype that couldn’t be maintained beyond two generations, suggesting some dominant *glh-1* alleles are too severe to recover with the current approach.

With the caveats of the GLH-1(D**Q**AD) in mind, significant increases in association with this mutant were primarily restricted to Argonaute proteins. These included not only the piRNA Argonaute PRG-1, but also the Argonautes CSR-1, WAGO-1, and C04F12.1, which bind to other small RNA species. In this regard, GLH-1(D**Q**AD) reflects the transient state of its insect and mammalian homologs that interact with piRNA amplifying Argonaute proteins but suggests the *C. elegans* transient complex is not limited to interactions with piRNAs. It is worth noting that the GLH-1 enriched proteins showing the most significant *decrease* in association with D**Q**AD are the PP2A subunits (PAA-1, PPTR-1, PPTR-2, and LET-92), whose phosphatase activity stabilizes P granules in the early embryo (Gallo et al., 2010; Griffin et al., 2011; Updike and Strome, 2009; Wang et al., 2014, Tables S2 and S3). This suggests that the targets of this phosphatase activity are not enriched in these D**Q**AD aggregates, and by extension may associate with GLH-1 in its open configuration but not this closed transient state.

Another exciting finding from this study is the enrichment of PCI complex “zomes” in the GLH-1 IPs. While GLH-1’s direct association with the COP9 signalosome component CSN5 was previously established, finding an enrichment for almost every PCI protein strongly suggests that GLH-1 has an affinity for these multilobed and structurally conserved scaffolding complexes. It will be imperative to understand how these scaffolds associate with GLH-1, the specific complex components that show direct interactions like CSN5, and if there is a spatial-temporal element to these interactions during germline development. Interestingly, while CSN5 is found in the cytoplasm and nucleus, it exhibits no distinct P-granule enrichment (Pintard et al., 2003; Smith et al., 2002), nor have other PCI subunits to date. One model is that COP9 and 26S Lid complex associate with GLH-1 in P granules, but another model is that they interact with GLH-1 that has found its way outside of P granules. eIF3 proteins, with the exception of eIF-3d, also show an affinity for GLH-1, while large and small ribosomal proteins show the inverse. As observed with CSN5, the enrichment of eIF3 subunits and the exclusion of ribosomal proteins in the GLH-1 IP does not necessarily equate with P-granule enrichment or exclusion; however, passive diffusion of proteins larger than 40kDa is prevented in P granules. While proteasome and ribosomal proteins smaller than 40kDA may not be excluded, P granules may act to compartmentalize germ plasm by excluding assembled 40S and 60S subunits and the 26S proteasome or its 20S core. Subsequent studies will investigate this possibility and directly address the impact of GLHs on protein turnover and translational regulation.

## STAR METHODS

### Strain Generation & Maintenance

*C. elegans* strains were maintained using standard protocols (Brenner, 1974). See Figure S2 for a complete list of GLH-1∷GFP∷3xFLAG alleles. Additional strains created for this study include DUP121 *glh-1(sam24[glh-1∷gfp∷3xFLAG]) I; pgl-1(sam52[pgl-1∷mTagRFPT∷3xFLAG]) IV*, DUP162 *glh-1(sam24[glh-1∷gfp∷3xFLAG]) I*; *itIs37[pie-1p∷mCherry∷H2B∷pie-1 3’UTR, unc-119(+)] IV*, DUP163 *glh-1(sam92[glh-1(D****Q****AD)∷gfp∷3xFLAG]) I*; *itIs37[pie-1p∷mCherry∷H2B∷pie-1 3’UTR, unc-119(+)] IV*, DUP165 *glh-2(sam82[glh-2(D****Q****AD)]) glh-1(sam92[glh-1(D****Q****AD)∷gfp∷3xFLAG]) I*; *itIs37[pie-1p∷mCherry∷H2B∷pie-1 3’UTR, unc-119(+)] IV*, DUP178 *glh-1(sam24[glh-1∷gfp∷3xFLAG]) prg-1(sam97[mTagRFP∷3xFLAG∷PRG-1]) I*, DUP180 *glh-1(sam65[Δglh-1∷gfp∷3xFLAG]) prg-1(sam97[TagRFP∷3xFLAG∷PRG-1]) I*, DUP181 *glh-1(sam92[glh-1(D****Q****AD)∷gfp∷3xFLAG]) prg-1(sam97[TagRFP∷3xFLAG∷PRG-1]) I*, and DUP184 *glh-1(sam86[glh-1(****_****EAD)∷gfp∷3xFLAG]) prg-1(sam97[TagRFP∷3xFLAG∷PRG-1]) I*. All strains generated for this study and their associated sequence files are available upon request.

### Screen Design

EMS mutagenesis was performed in DUP64 *glh-1(sam24[glh-1∷GFP∷3xFLAG]) I* worms using the standard protocol (Kutscher and Shaham, 2014). Adult grandchildren (F2s) were then screened under a Leica M165FC fluorescence stereomicroscope for changes in P-granule appearance as visualized with the GLH-1∷GFP∷3xFLAG reporter. Sequencing of *glh-1* was performed for several mutants exhibiting aberrant GLH-1∷GFP phenotypes.

### CRISPR strain construction

Creation of the *glh-1(sam24[glh-1∷GFP∷3xFLAG]) I* allele was previously described (Andralojc et al., 2017). A co-CRISPR technique with *dpy-*10 was used to create the mutant alleles as described (Paix et al., 2017). Table S1 lists the sequences for the guide RNA and repair templates for the strains created. An mTagRFPT∷3xFLAG tag was added to the C-terminus of *pgl-1* and the N-terminus of *prg-1* using the FP-SEC method to create *pgl-1*(*sam52)* and *prg-1(sam97)* alleles (Dickinson et al., 2015). The same method was modified to generate the *glh-1(sam31)* allele found in DUP73, which contains a bicistronic rSL2 GFP∷3xFLAG transcriptional *glh-1* reporter. All edits generated for this study were sequence verified, and sequence and genebank files for each strain are available upon request.

### Fertility Assay

For each strain the fertility was determined by plating L4 worms at both 20°C and 26°C. Hatched F1 progeny were then picked to 10 plates with 25 worms on each plate. The percent of grotty (uterus filled with unfertilized oocytes and terminal embryos) and clean (germline atrophy with empty uterus) sterile F1s were scored when they reached day 2 of adulthood.

### Embryonic Lethality

For each strain embryonic lethality was determined by plating L4 worms at both 20°C and 26°C. Hatched F1s were grown to adulthood, and six worms were picked to new plates and allowed to lay for five hours. Embryos were marked and counted, and unhatched embryos were counted again 18-24 hours later. Terminal phenotypes were imaged from these unhatched, but still moving, embryos.

### Imaging, P-granule counts, and expression intensity

Live worms were mounted on agarose pads in egg buffer (25 mM HEPES, 120 mM NaCl, 2 mM MgCl2, 2 mM CaCl2, 50 mM KCl, and 1 mM levamisole) and imaged with a 63X objective under fixed exposure conditions. A single plane focused mid-way through the germline loop region was acquired for ten worms from each strain using Leica AF6000 acquisition software on an inverted Leica DMI6000B microscope with an attached Leica DFC365FX camera. An ImageJ was used to count P granules and average GFP intensity within the field of view.

To measure P-granule size in P4 cells (Figure 5), live embryos from DUP162, DUP163, and DUP165 were mounted on agarose slides and focused mid-way through P4 nuclei using the mCherry∷H2B signal. Both mCh and GFP were acquired with fixed exposure conditions at this single focal plane. ImageJ was used to identify and calculate P-granule area in at least 40 embryos from each strain.

For higher resolution images in Figure 1A, 2C, 4, and 5, the same microscope and objective was used to acquire Z sections through 10 microns, and then images were deconvolved and shown as a max intensity projection. All scale bars are 20 microns. For immunostained images (Figure 4A/B/C), worms and embryos were fixed and stained with anti-GFP monoclonal and anti-PGL-1 polyclonal antibodies as previously described (Andralojc et al., 2017).

### Taqman Assay to quantify 21UR-1 abundance

21UR-1 abundance was quantified relative to miR-1 using a TaqMan-based small RNA qPCR reaction following established protocols (Han et al., 2009). 21UR-1 (TGGTACGTACGTTAACCGTGC) and miR-1 (TGGAATGTAAAGAAGTATGTA) primers were previously validated (Montgomery et al., 2012). Averages and the standard deviation from five to six replicates of each strain are shown in Figure S3a. RNA was prepared from a fixed number of young adult worms with healthy, GFP-expressing germlines.

### Liquid Culture

25 plates from each of the DUP64, DUP73, DUP171 and DUP 173 strains were grown until gravid and then bleach treated to harvest embryos. Hatched embryos were used to seed 250 ul of S Media (separated into four 1 liter flasks) containing freeze dried OP50 (LabTie) on the following day. Worms were grown in liquid culture and until most reached the young adult stage with embryos in the uterus, and then harvested, washed, and bleach treated to synchronize development. L1-stage worms hatched without food were then used to inoculate another 250 ul of S Media with OP50. Worms were grown until the majority reached the young adult stage, where they were harvested, washed, and flash frozen in 1ml aliquots.

### Preparation of protein lysate

1-2 mL of frozen worms grown from liquid culture were removed from the −80 and ground with a mortar and pestle in liquid nitrogen for 15 minutes, and then resuspended in 7 mL of cold lysis buffer (25mM Hepes-KOH (pH 7.4), 10mM KOAc, 2mM Mg(OAc)^2^, 100mM KCl, 0.25% Triton X-100, 1mM PMSF, 1 proteinase inhibitor tablet, 1:200 Riboguard and 1:200 DNase). Samples were then spun at 5,000 RPM for 5 minutes and supernatant was then immediately used for immunoprecipitation.

### Immunoprecipitation

Anti-DYKDDDDK agarose beads (018-22783, Wako Chemicals, USA) were removed from the −20 and allowed to equilibrate for 10 mins and were then inverted to resuspend the beads. 100 μL of the bead suspension was then aliquoted into 7 Eppendorf tubes. 1 mL of cold 1x TBS was added to each tube. Beads were then spun at 4C for 1 min at 1200 RPMs. Supernatant was removed and beads were washed with 1 mL of cold 1x TBS and then centrifuged in the 4C for 1 min at 1200 RPMs. The supernatant was removed with a pipette and then the TBS wash was repeated two more times. After the last wash 1 mL of the protein lysate was pipetted into each of the 7 Eppendorf tubes with agarose beads. Tubes were inverted to mix and then placed on a rotator in the 4C for 3 hours. After 3 hours all the tubes were spun in the 4C for 1 min at 1200 RPMs. The supernatant, which contained the unbound fraction, was removed and then beads were washed and spun 3 times with 1 mL of 1x TBS. During the 3^rd^ wash 10 μL was removed and imaged to confirm the GFP tagged protein was attached to the beads (see imaging agarose beads section). Beads from all the 7 tubes were combined into 1 Eppendorf tube and flash frozen in liquid nitrogen. Beads with protein still bound were then sent to Cell Signaling Technologies for Mass spectrometry.

15ul of anti-DYKDDDDK agarose beads under control or protein exposed conditions were imaged under DIC and GFP with a 10X objective on the Leica DMI6000B microscope (Figure S3). A western was used to demonstrate that conditions were optimal for GLH-1∷GFP∷3xFLAG immunoprecipitation.

### Mass Spectrometry and Analysis

The mass spectrometry was performed by Cell Signaling Technologies (CST) (Danvers, MA) using their PTMScan Discovery service. On bead protease digestion was performed followed by C18 solid phase extraction, peptide lyophilization, and antibody enrichment of post-translational modification (PTM) containing peptides using CST developed antibodies. Then peptides were loaded onto a 50 cm x 100 μm PicoFrit capillary column packed with C18 reversed-phase resin for LC-MS/MS. The MS parameters were as follows: MS Run Time 168 min, MS1 Scan Range (300.0 – 1500.00), Top 20 MS/MS (Min Signal 500, Isolation Width 2.0, Normalized Coll. Energy 35.0, Activation-Q 0.250, Activation Time 20.0, Lock Mass 371.101237, Charge State Rejection Enabled, Charge State 1+ Rejected, Dynamic Exclusion Enabled, Repeat Count 1, Repeat Duration 35.0, Exclusion List Size 500, Exclusion Duration 40.0, Exclusion Mass Width Relative to Mass, Exclusion Mass Width 10ppm). Mass spectrometry informatics was also done by Cell Signaling Technologies. The MS/MS spectra were evaluated using SEQUEST (Eng et al., 1994) and the Core platform from Harvard University. Searches were performed against the most recent update of the WormBase *C elegans* database with mass accuracy of +/-50 ppm for precursor ions and 0.02 Da for product ions. Results were then filtered with mass accuracy of +/– 5 ppm on precursor ions and presence of the intended motif. These were then further filtered to a 1% protein false discovery rate. Analysis results are included in Table S2, which has also been uploaded to the PRIDE proteomics repository, project accession PXD014135.

## ACKNOWLEDGEMENTS

We would like to thank Chris Smith in the MDI Biological Laboratory Sequencing Core for her assistance in sequencing and genotyping strains generated for this publication, Dorothy Wheatcraft at the Mass Spectrometry facility at the Jackson Laboratory for proteomics assistance and advice, Matt Stokes at Cell Signaling Technology, Inc. for performing the proteomics and analysis of our GLH-1 IPs, and Tai Montgomery at Colorado State University for providing protocols and primer sets to quantify piRNA abundance in our mutants.

The Updike lab is supported by NIH-NIGMS [R01GM113933]. E.A.M. is supported by a postdoctoral NRSA fellowship NIH-NIGMS [F32GM128248]. Equipment and cores used for parts of this study are supported by NIH-NIGMS COBRE [P20GM104318] and INBRE [P20GM203423]. Undergraduate support included an NSF-REU [DBI-1460495] fellowship to J.H.F. and MDIBL’s James Slater Murphy fellowship to E.L.X.

---

<u>Figure S1.</u> Comparison of the amino acid sequence conservation between *C. elegans* DEAD-box helicase proteins, drosophila Vasa and human DDX4 in A) the flanking domain B) the negatively charged C terminal tryptophan domain. C) The phylogenomic tree showing the conservation of the CCHC zinc-knuckle motif (orange) across Vasa proteins has been updated from (Figure 1C in Gustafson and Wessel, 2010) using phyloT (phylot.biobyte.de) and NCBI taxonomy data. Dashed lines indicate loss of the motif in some species.

<u>Figure S2.</u> All images (now condensed) used to quantify granularity and expression in Figure 3. The domain location of the mutation, strain name, allele name and mutation type is indicated.

<u>Figure S3.</u> A) TaqMan-based small-RNA qRT-PCR comparing abundance of the 21UR-1 piRNA relative to miR-1. B) GLH-1 Immunoprecipitation. Top: Microscope images of control non-antigen exposed anti-DYKDDDDK agarose beads compared to beads exposed to lysate from the GLH-1∷GFP∷3xFLAG expressing strain. DIC (left), GFP (right). Bottom: Western blot comparing GLH-1 expression in the input, unbound and elute fraction after immunoprecipitation with protein lysate from the GLH-1∷GFP∷3xFLAG expressing strain.

Table S2: Proteins with enhanced or depleted GLH-1 association in wild type (DEAD) and GLH-1 mutants (DAAD & DQAD) 2505 proteins identified by FLAG-IP LC/MS/MS of 3xFLAG-tagged GLH-1 (and mutant GLH-1) *C. elegans* lysates compared to control. Large Excel table (not attached). This data has been uploaded to the PRIDE proteomics repository, project accession PXD014135.

**Table S1:**
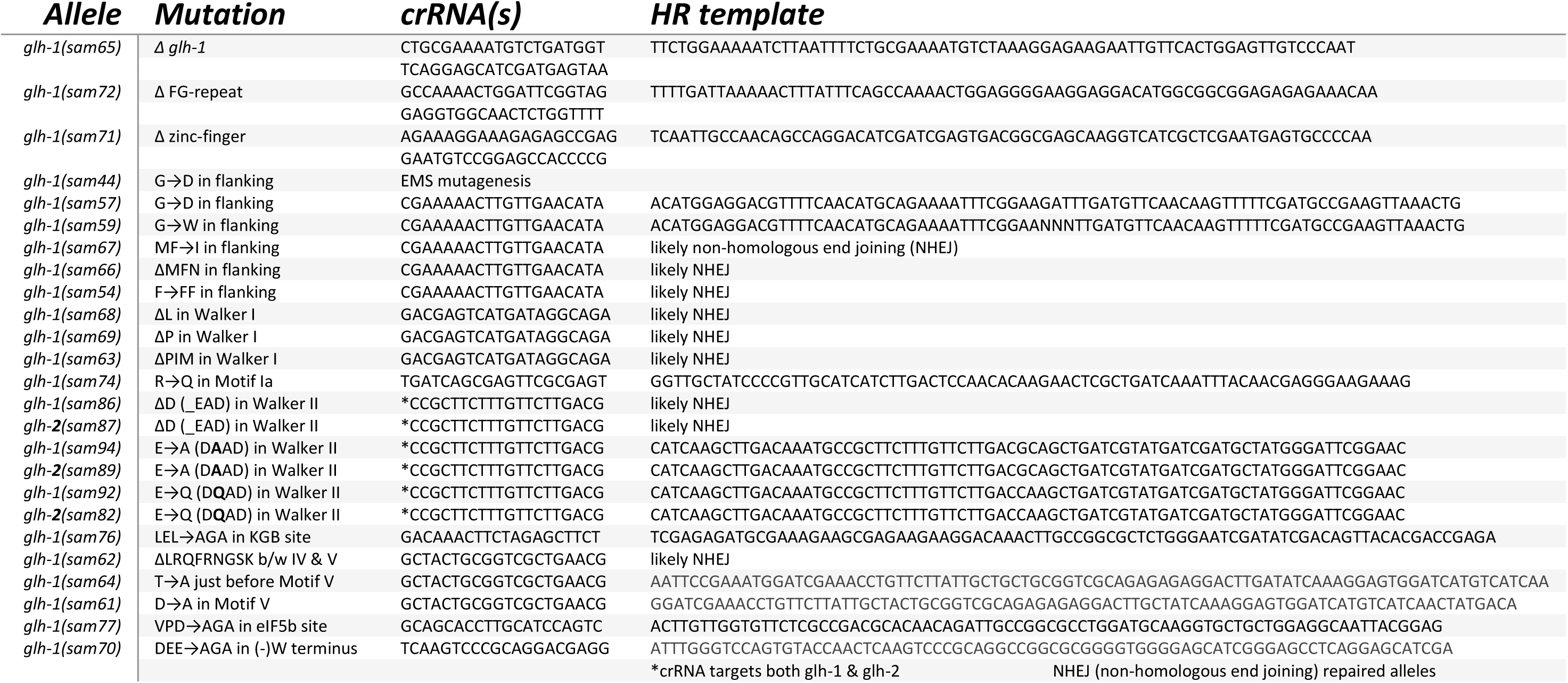
CRISPR/Cas9 reagents for generating *glh-1* alleles.

**Table S3:**
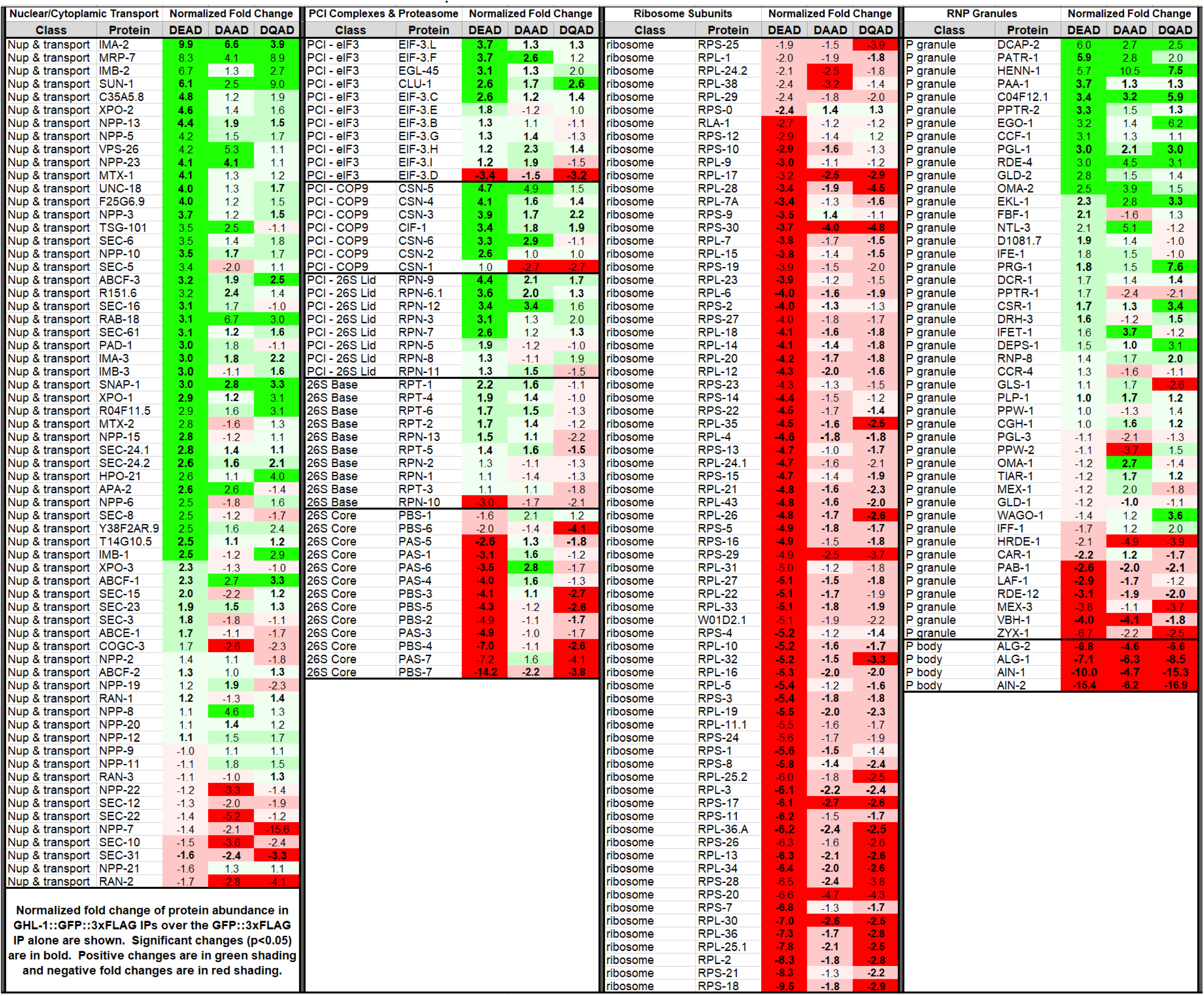
Protein classes with enriched or depleted GLH-1 associations.

