## Supplementary figures and images for "Germline maintenance through the multifaceted activities of GLH/Vasa in *Caenorhabditis elegans* P granules"

### Figure S1

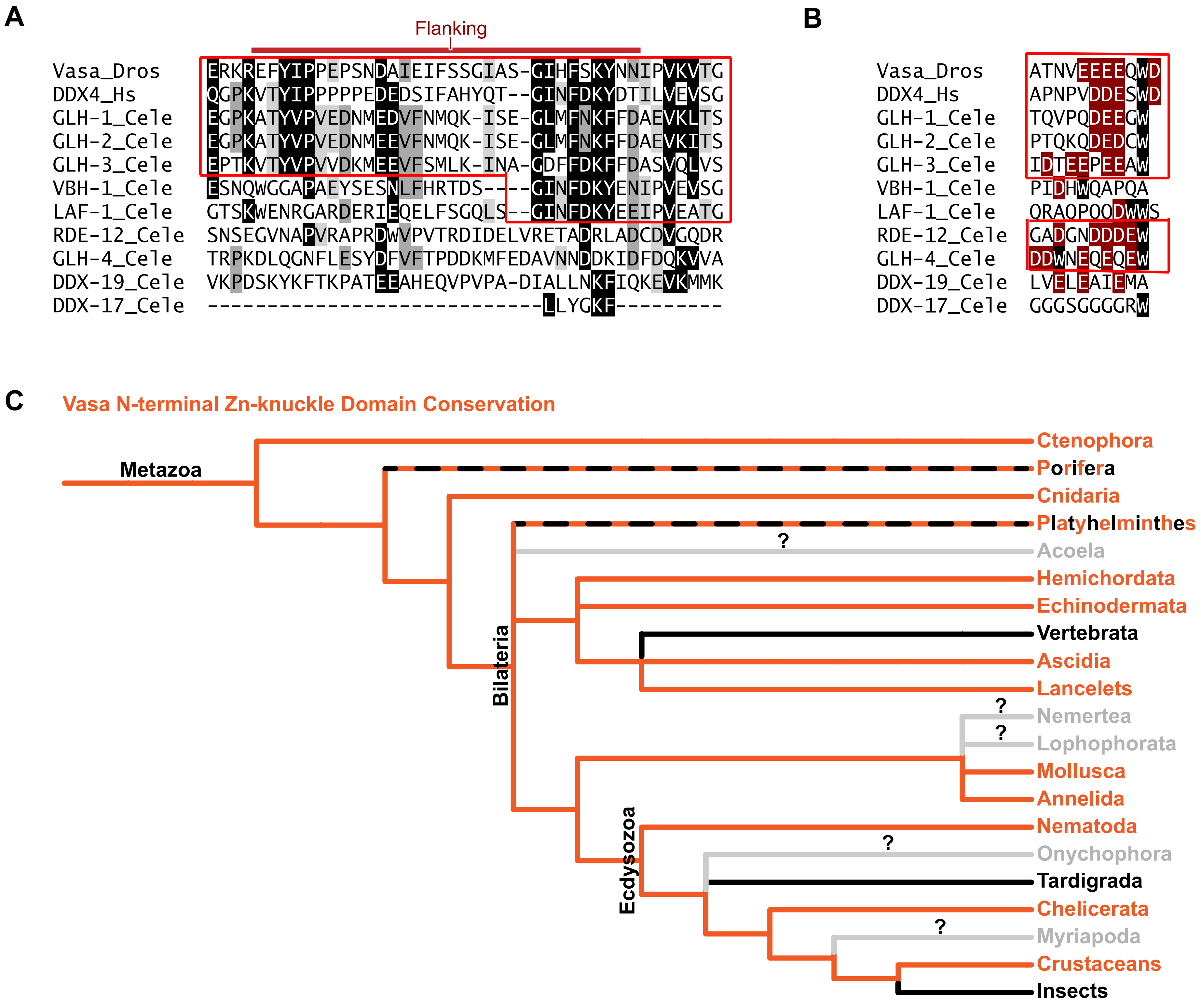

### Figure S3

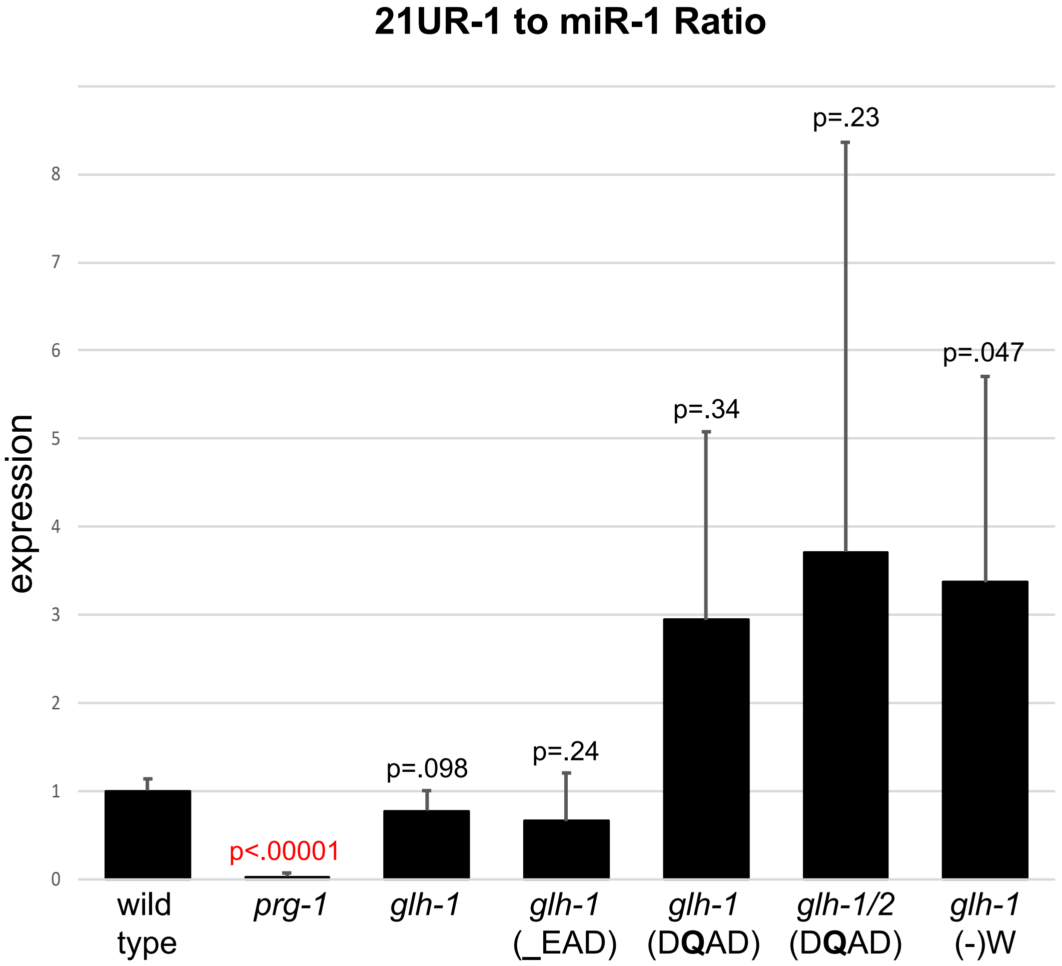

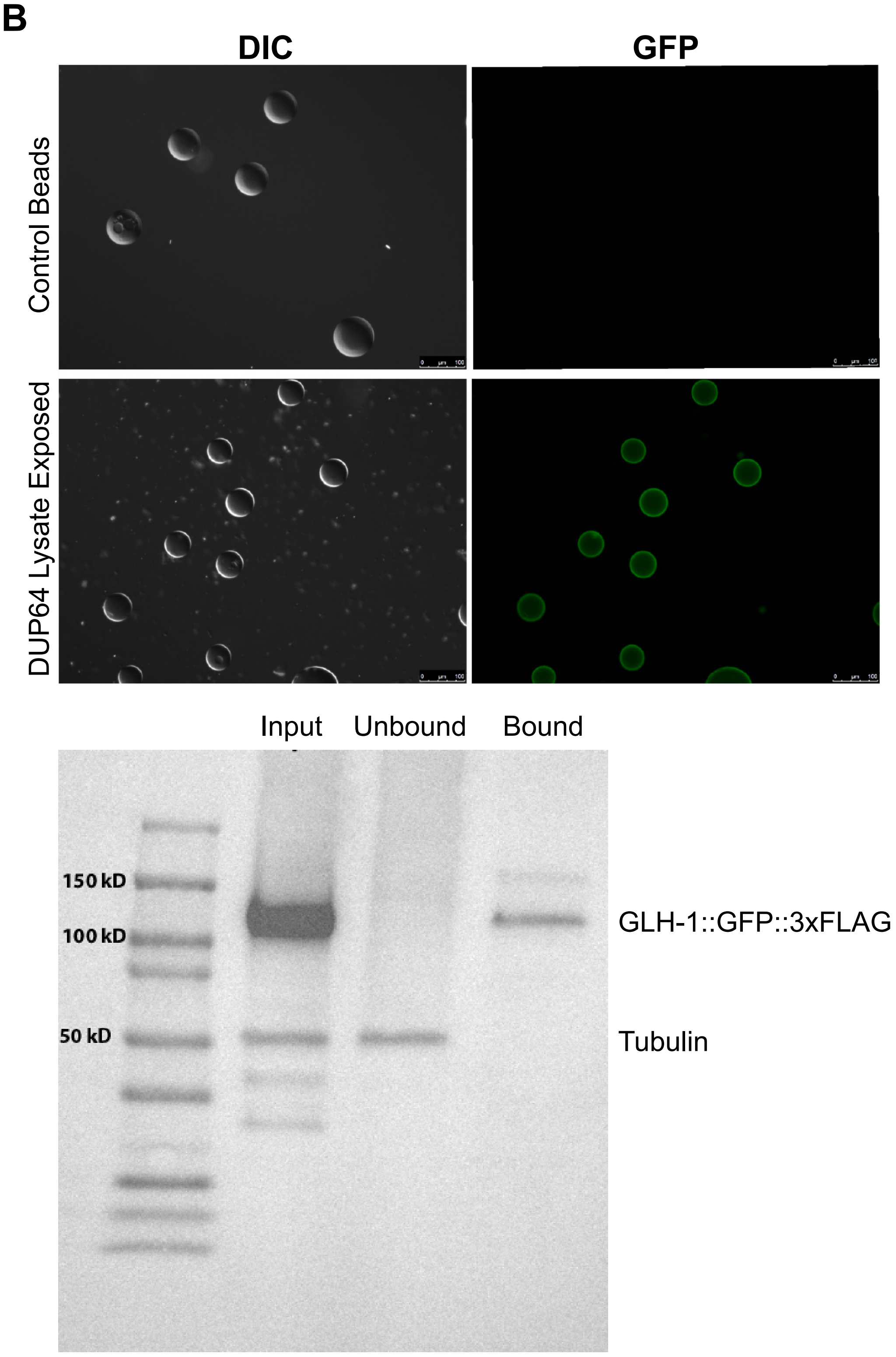
