## Supplementary material for "Germline maintenance through the multifaceted activities of GLH/Vasa in *Caenorhabditis elegans* P granules": Figure S2

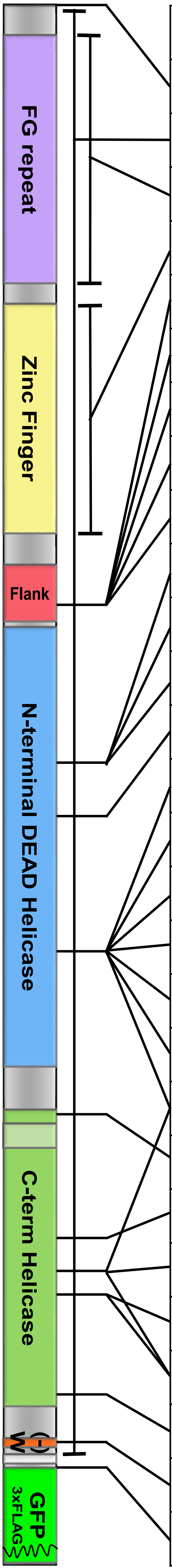


| Strain | Allele | Mutation |
|---|---|---|
| DUP64 | *sam24* | GLH-1::GFP (parental strain) |
| DUP144 | *sam65* | Δ*glh-1* (*glh-1* deletion) |
| DUP152 | *sam72* | ΔFG (FG repeat deletion) |
| DUP151 | *sam71* | ΔZF (zinc finger deletion) |
| DUP130 | *sam57* | G→D |
| DUP137 | *sam59* | G→W |
| DUP146 | *sam67* | MF→I |
| DUP145 | *sam66* | ΔMFN |
| DUP124 | *sam54* | F→FF |
| DUP147 | *sam68* | ΔL |
| DUP148 | *sam69* | ΔP |
| DUP142 | *sam63* | ΔPIM |
| DUP154 | *sam74* | R→Q |
| DUP168 | *sam86* | ΔD in *glh-1* only (_EAD) |
| DUP175 | *sam87* *sam86* | ΔD in *glh-1* & *glh-2* (_EAD) |
| DUP173 | *sam94* | E→A in *glh-1* only (DAAD) |
| DUP170 | *sam89* *sam94* | E→A in *glh-1* & *glh-2* (DAAD) |
| DUP171 | *sam92* | E→Q in *glh-1* only (DQAD) |
| DUP160 | *sam82* *sam92* | E→Q in *glh-1* & *glh-2* (DQAD) |
| DUP179 | *sam92* *sam64* | E→Q (DQAD) + T→A (no ATPase) |
| DUP156 | *sam76* | LEL→AGA |
| DUP141 | *sam62* | ΔLRQFRNGSK |
| DUP143 | *sam64* | T→A (no ATPase) |
| DUP140 | *sam61* | D→A |
| DUP164 | *sam61* *sam64* | D→A + T→A (no ATPase) |
| DUP157 | *sam77* | VPD→AGA |
| DUP149 | *sam70* | DEE→AGA |
| DUP73 | *sam31* | *glh-1* txn reporter (bicistronic GFP) |

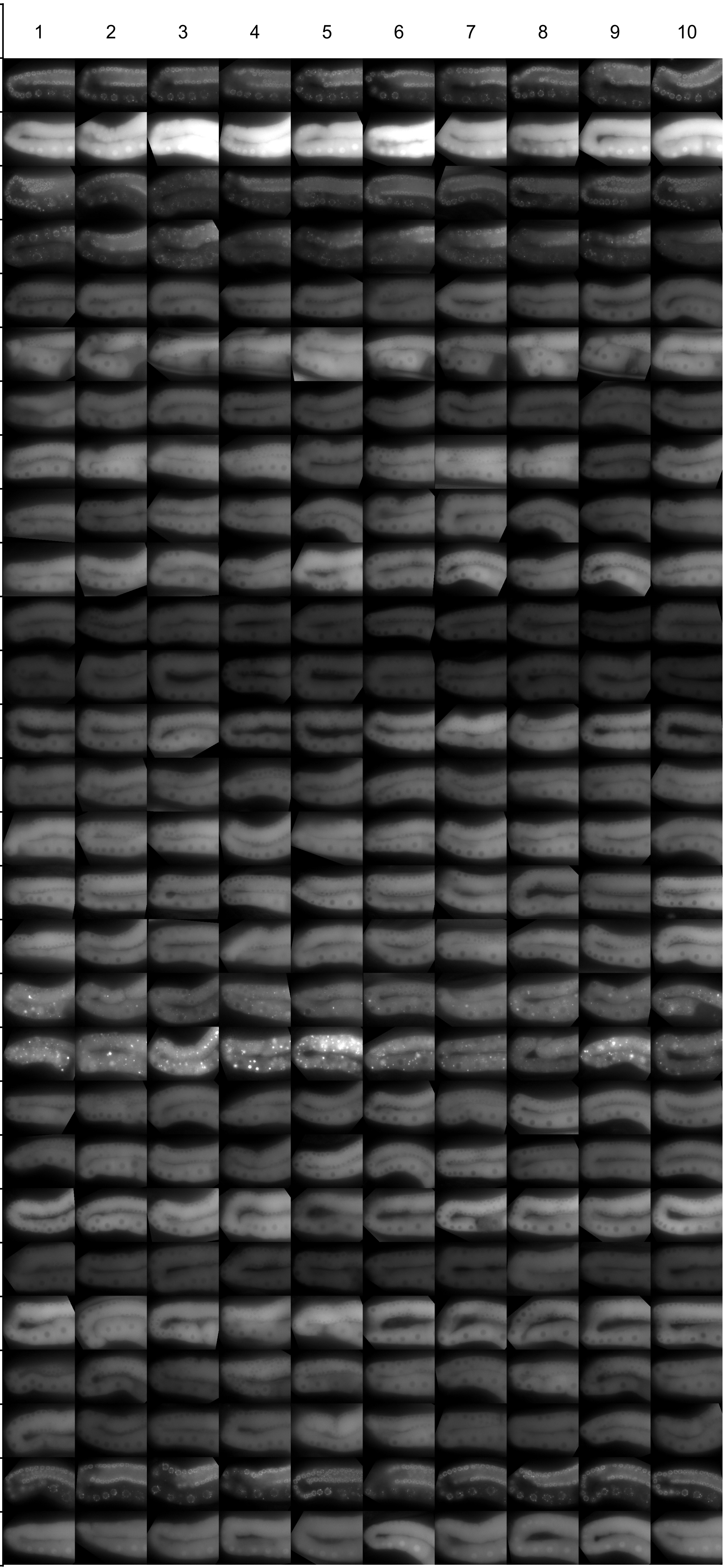
