## Supplementary material for "Germline maintenance through the multifaceted activities of GLH/Vasa in *Caenorhabditis elegans* P granules": Table S1

| **Allele** | **Mutation** | **crRNA(s)** | **HR template** |
| --- | --- | --- | --- |
| glh-1(sam65) | *Δ glh-1* | CTGCGAAAATGTCTGATGGT | TTCTGGAAAAATCTTAATTTTCTGCGAAAATGTCTAAAGGAGAAGAATTGTTCACTGGAGTTGTCCCAAT |
|  |  | TCAGGAGCATCGATGAGTAA |  |
| glh-1(sam72) | Δ FG-repeat | GCCAAAACTGGATTCGGTAG | TTTTGATTAAAAACTTTATTTCAGCCAAAACTGGAGGGGAAGGAGGACATGGCGGCGGAGAGAGAAACAA |
|  |  | GAGGTGGCAACTCTGGTTTT |  |
| glh-1(sam71) | Δ zinc-finger | AGAAAGGAAAGAGAGCCGAG | TCAATTGCCAACAGCCAGGACATCGATCGAGTGACGGCGAGCAAGGTCATCGCTCGAATGAGTGCCCCAA |
|  |  | GAATGTCCGGAGCCACCCCG |  |
| glh-1(sam44) | G→D in flanking | EMS mutagenesis |  |
| glh-1(sam57) | G→D in flanking | CGAAAAACTTGTTGAACATA | ACATGGAGGACGTTTTCAACATGCAGAAAATTTCGGAAGATTTGATGTTCAACAAGTTTTTCGATGCCGAAGTTAAACTG |
| glh-1(sam59) | G→W in flanking | CGAAAAACTTGTTGAACATA | ACATGGAGGACGTTTTCAACATGCAGAAAATTTCGGAANNNTTGATGTTCAACAAGTTTTTCGATGCCGAAGTTAAACTG |
| glh-1(sam67) | MF→I in flanking | CGAAAAACTTGTTGAACATA | likely non-homologous end joining (NHEJ) |
| glh-1(sam66) | ΔMFN in flanking | CGAAAAACTTGTTGAACATA | likely NHEJ |
| glh-1(sam54) | F→FF in flanking | CGAAAAACTTGTTGAACATA | likely NHEJ |
| glh-1(sam68) | ΔL in Walker I | GACGAGTCATGATAGGCAGA | likely NHEJ |
| glh-1(sam69) | ΔP in Walker I | GACGAGTCATGATAGGCAGA | likely NHEJ |
| glh-1(sam63) | ΔPIM in Walker I | GACGAGTCATGATAGGCAGA | likely NHEJ |
| glh-1(sam74) | R→Q in Motif Ia | TGATCAGCGAGTTCGCGAGT | GGTTGCTATCCCCGTTGCATCATCTTGACTCCAACACAAGAACTCGCTGATCAAATTTACAACGAGGGAAGAAAG |
| glh-1(sam86) | ΔD (_EAD) in Walker II | *CCGCTTCTTTGTTCTTGACG | likely NHEJ |
| glh-**2**(sam87) | ΔD (_EAD) in Walker II | *CCGCTTCTTTGTTCTTGACG | likely NHEJ |
| glh-1(sam94) | E→A (D**A**AD) in Walker II | *CCGCTTCTTTGTTCTTGACG | CATCAAGCTTGACAAATGCCGCTTCTTTGTTCTTGACGCAGCTGATCGTATGATCGATGCTATGGGATTCGGAAC |
| glh-**2**(sam89) | E→A (D**A**AD) in Walker II | *CCGCTTCTTTGTTCTTGACG | CATCAAGCTTGACAAATGCCGCTTCTTTGTTCTTGACGCAGCTGATCGTATGATCGATGCTATGGGATTCGGAAC |
| glh-1(sam92) | E→Q (D**Q**AD) in Walker II | *CCGCTTCTTTGTTCTTGACG | CATCAAGCTTGACAAATGCCGCTTCTTTGTTCTTGACCAAGCTGATCGTATGATCGATGCTATGGGATTCGGAAC |
| glh-**2**(sam82) | E→Q (D**Q**AD) in Walker II | *CCGCTTCTTTGTTCTTGACG | CATCAAGCTTGACAAATGCCGCTTCTTTGTTCTTGACCAAGCTGATCGTATGATCGATGCTATGGGATTCGGAAC |
| glh-1(sam76) | LEL→AGA in KGB site | GACAAACTTCTAGAGCTTCT | TCGAGAGATGCGAAAGAAGCGAGAAGAAGGACAAACTTGCCGGCGCTCTGGGAATCGATATCGACAGTTACACGACCGAGA |
| glh-1(sam62) | ΔLRQFRNGSK b/w IV & V | GCTACTGCGGTCGCTGAACG | likely NHEJ |
| glh-1(sam64) | T→A just before Motif V | GCTACTGCGGTCGCTGAACG | AATTCCGAAATGGATCGAAACCTGTTCTTATTGCTGCTGCGGTCGCAGAGAGAGGACTTGATATCAAAGGAGTGGATCATGTCATCAA |
| glh-1(sam61) | D→A in Motif V | GCTACTGCGGTCGCTGAACG | GGATCGAAACCTGTTCTTATTGCTACTGCGGTCGCAGAGAGAGGACTTGCTATCAAAGGAGTGGATCATGTCATCAACTATGACA |
| glh-1(sam77) | VPD→AGA in eIF5b site | GCAGCACCTTGCATCCAGTC | ACTTGTTGGTGTTCTCGCCGACGCACAACAGATTGCCGGCGCCTGGATGCAAGGTGCTGCTGGAGGCAATTACGGAG |
| glh-1(sam70) | DEE→AGA in (-)W terminus | TCAAGTCCCGCAGGACGAGG | ATTTGGGTCCAGTGTACCAACTCAAGTCCCGCAGGCCGGCGCGGGGTGGGGAGCATCGGGAGCCTCAGGAGCATCGA |
|  |  |  | *crRNA targets both glh-1 & glh-2 |
